# Leveraging Heterogeneous Network Embedding for Metabolic Pathway Prediction

**DOI:** 10.1101/2020.02.20.940205

**Authors:** Abdur Rahman M. A. Basher, Steven J. Hallam

## Abstract

Metabolic pathway reconstruction from genomic sequence information is a key step in predicting regulatory and functional potential of cells at the individual, population and community levels of organization. Although the most common methods for metabolic pathway reconstruction are gene-centric e.g. mapping annotated proteins onto known pathways using a reference database, pathway-centric methods based on heuristics or machine learning to infer pathway presence provide a powerful engine for hypothesis generation in biological systems. Such methods rely on rule sets or rich feature information that may not be known or readily accessible. Here, we present pathway2vec, a software package consisting of six representational learning based modules used to automatically generate features for pathway inference. Specifically, we build a three layered network composed of compounds, enzymes, and pathways, where nodes within a layer manifest inter-interactions and nodes between layers manifest betweenness interactions. This layered architecture captures relevant relationships used to learn a neural embedding-based low-dimensional space of metabolic features. We benchmark pathway2vec performance based on node-clustering, embedding visualization and pathway prediction using MetaCyc as a trusted source. In the pathway prediction task, results indicate that it is possible to leverage embeddings to improve pathway prediction outcomes.

**Availability and implementation:** The software package, and installation instructions are published on github.com/pathway2vec

## 1 Introduction

Metabolic pathway reconstruction from genomic sequence information is a key step in predicting regulatory and functional potential of cells at the individual, population and community levels of organization. ([3]). Exponential advances in sequencing throughput continue to lower the cost of data generation with concomitant increases in data volume and complexity ([4]). Resulting data sets create new opportunities for metabolic reconstruction within biological systems that require the development of new computational tools and approaches that scale with data volume and complexity. Although the most common methods for metabolic pathway reconstruction are gene-centric e.g. mapping annotated proteins onto known pathways using a reference database based on sequence homology, heuristic or rule-based methods for pathway-centric inference including PathoLogic ([21]) and MinPath ([35]) have become increasingly used to generate hypotheses and build quantitative models. For example, Pathologic generates pathway genome databases (PGDBs) that can be refined based on experimental validation e.g. EcoCyc ([22]) and stored in repositories e.g. BioCyc ([9]) for community access and use in flux balance analysis.

The development of accurate and flexible rule sets for pathway prediction remains a challenging enterprise informed by expert curators incorporating thermodynamic, kinetic, and structural information for validation ([33]). Updating these rule sets as new organisms or pathways are described and validated can be cumbersome and out of phase with current user needs. This has led to the consideration of machine learning (ML) approaches for pathway prediction based on rich feature information. Dale and colleagues conducted a seminal study comparing the performance of Pathologic to different types of supervised ML algorithms (naive Bayes, k nearest neighbors, decision trees and logistic regression), converting rules into features, defining new features, and evaluating on experimentally validated pathways from six highly curated organisms in the BioCyc collection randomly divided into training and test sets ([11]). Resulting performance metrics indicated that generic ML methods equaled and in some cases exceeded performance of Pathologic with the benefit of probability estimation for pathway presence and increased flexibility and transparency of use.

Despite the potential benefits of adopting ML methods for pathway prediction from genomic sequence information, Pathologic remains the primary inference engine of Pathway Tools ([21]), and alternative methods for pathway-centric inference expanding on the generic methods described above remain nascent. Several of these methods incorporate metabolite information to improve pathway inference and reaction rules to infer metabolic pathways ([7, 33, 32]). Other methods including BiomeNet ([29]) and MetaNetSim ([19]) dispense with pathways all together and model reaction networks based on enzyme abundance information. Recently, Basher et. al ([6]) implemented a multi-label classification approach to predict metabolic pathways for individual genomes as well as more complex cellular communities e.g. microbiomes. One of the primary challenges encountered in developing mlLGPR relates to engineering reliable features representing heterogeneous and degenerate functions within multi-organism data sets ([23]).

Advances in representational learning have led to the development of scalable methods for engineering features from graphical networks e.g. networks composed of multiple nodes including information systems or social networks ([16, 12, 28])). These approaches learn feature vectors for nodes in a network by solving an optimization problem in an unsupervised manner, using random walks followed by Skip-Gram extraction of low dimensional latent continuous features, known as *embeddings* ([25]). Here we present pathway2vec, a software package incorporating multiple *random walks based* algorithms for representational learning used to automatically generate feature representations of metabolic pathways, which are decomposed into three interacting layers: compounds, enzymes and pathways, where each layer consists of associated nodes. A Skip-Gram model is applied to extract embeddings for each node, encoding smooth decision boundaries between groups of nodes in that graph. Nodes within a layer manifest inter-interactions and nodes between layers manifest betweenness interactions resulting in a multi-layer heterogeneous information network ([30]). This layered architecture captures relevant relationships used to learn a neural embedding-based low-dimensional space of metabolic features (Fig. 1).

**Figure 1:**
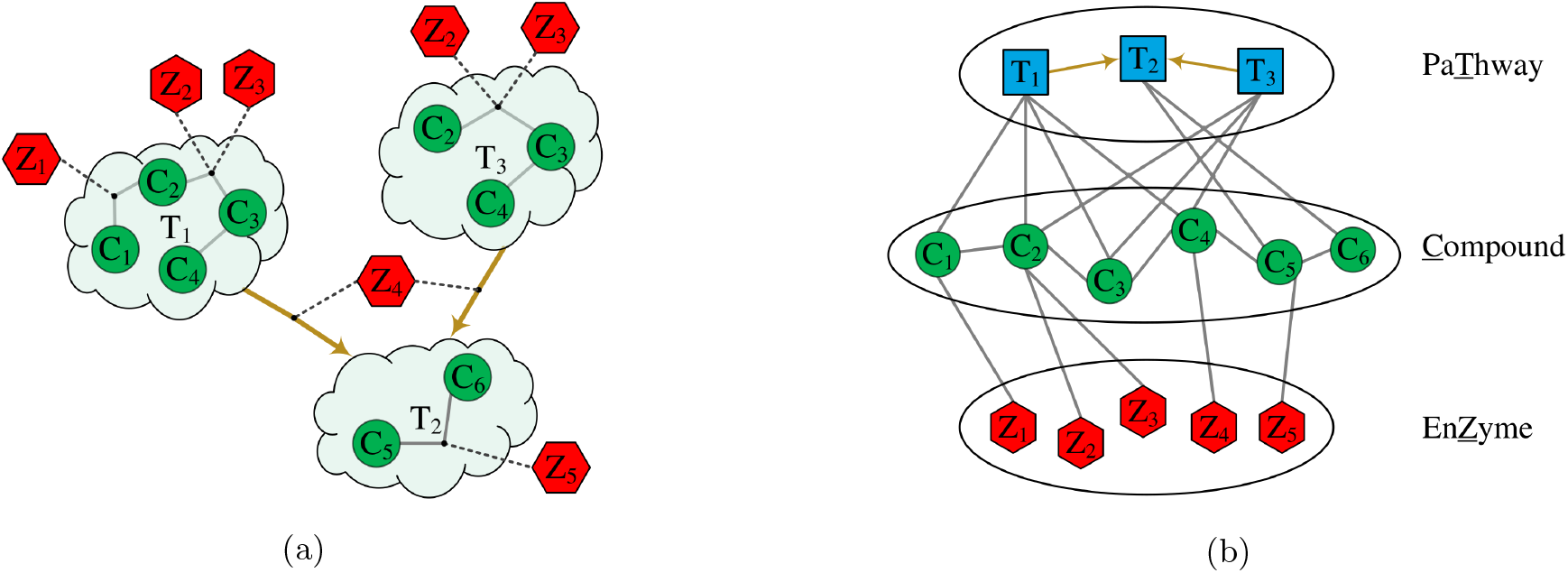
Three interacting metabolic pathways (a), depicted as a cloud glyph, where each pathway is comprised of compounds (green) and enzymes (red). Interacting compound, enzyme and pathway components are transformed into a multi-layer heterogeneous information network (b).

In addition to implementing several published random walk methods, we developed RUST (unit-ci<u>r</u>cle based j<u>u</u>mp and <u>st</u>ay random walk), adopting a unit-circle equation to sample node pairs that generalize previous random walk methods ([16, 12, 18]). The modules in pathway2vec were benchmarked based on node-clustering, embedding visualization, and pathway prediction. In the case of pathway prediction, pathway2vec modules provided a viable adjunct or alternative to manually curated feature sets used in ML based metabolic pathway reconstruction from genomic sequence information. The distinctness of this work lies in decomposing pathway into components, so various graph learning methods can be applied to automatically extract semantic features of metabolic pathways, and to incorporate the learned embeddings for pathway inference.

## 2 Definitions and Problem Statement

In this section, we formulate the problem of metabolic features engineering using a heterogeneous information network. Throughout the paper, all vectors are column vectors denoted by boldface lowercase letters (e.g., x) while matrices are represented by boldface uppercase letters (e.g., **X**). The **X**_*i*_ matrix denotes the *i*-th row of **X** and **X***_i,j_* denotes the (*i*,*j*)-th element of **X**. A subscript character to a vector, x_*i*_, denotes an *i*-th cell of x. Occasional superscript, **X**^(*i*)^, suggests an index to a sample, position, or current epoch during learning period. We use calligraphic letters to represent sets (e.g., 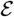) while we use the notation |.| to denote the cardinality of a given set.

### Definition 2.1. Multi-label Pathway Dataset ([6])

A pathway dataset is characterized by 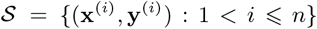 consisting of *n* examples, where x^(*i*)^ is a vector indicating abundance information for each enzymatic reaction denoted by *z*, which is an element of a set 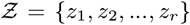, having *r* possible reactions. The abundance of an enzymatic reaction for a given example *i*, say 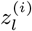, is defined as 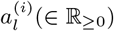. The class label 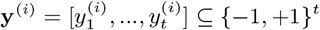 is a pathway label vector of size *t* representing the total number of pathways obtained from a trusted source of experimentally validated metabolic pathways 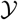. The matrix form of x^(*i*)^ and y^(*i*)^ are symbolized as **X** and **Y**, respectively. ■

Both 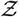 and 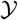 are derived from trusted sources, such as KEGG ([20]) or MetaCyc ([8]). We assume that there is a numerical representation behind every instance and label.

The pathway inference task can be formulated as retrieving a set of pathway labels for an example *i* given features learned according to a *heterogeneous information network* defined as:

### Definition 2.2. Heterogeneous Information Network

A heterogeneous information network is defined as a graph 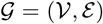, where 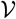 and 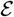 denote to the set of nodes and edges (either directed or undirected), respectively ([31]). Each 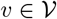 is associated with an object type mapping function 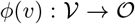, where 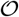 represents a set of object types. Each edge 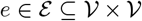 includes multiple types of links, and is associated with a link type mapping function 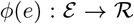, where 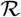 represents a set of relation types. In particular, when 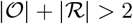, the graph is referred to as a heterogeneous information network. ■

In heterogeneous information networks, both object types and relationship types are explicitly segregated. For the undirected edges, notice that if a relation exists from a type 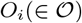 to a type 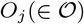, denoted as *O_i_RO_j_* and 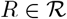, the inverse relation *R*^−1^ holds naturally for *O_j_R*^−1^*O_i_*. However, in many circumstances, *R* and its inverse *R*^−1^ are not equal, unless the two objects are in the same domain, and *R* is symmetric. In addition, the network may be weighted where each edge *e_i,j_*, of nodes *i* and *j*, is associated with a weight of type *R*. The linkage type of an edge automatically defines the node types of it’s end points. The graph articulated in this paper is considered directed and weighted (in some cases), but for simplification is converted to a undirected network by simply treating edges as symmetric links. Note that if 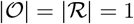, the network is *homogeneous*; otherwise, it is heterogeneous.

### Example 2.2.1.

MetaCyc can be abstracted as a heterogeneous information network, in Fig. 1(b), which contains 3 types of objects, namely compounds (C), enzymes (Z), and pathways (T). There exist different types of links between objects representing semantic relationships e.g. “*composed of*” and “*involved in*”, relationships between pathways and compounds or relations between enzymes and compounds e.g. “*transform*” and “*transformed by*”. An enzyme may be mapped to a numerical category, known as an enzyme commission number (EC) based on the chemical reaction it catalyzes.

Two objects within heterogeneous information networks describe meta-level relationships refereed to as *meta-paths* ([31]).

### Definition 2.3. Meta-Path

A meta-path 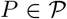 is a path over 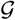 in the form of 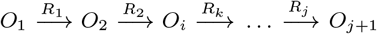, which defines an aggregation of relationships *U* = *R*_1_ ∘ *R*_2_ ∘ … ∘ *R_j_* between type *O*_1_ and *O*_*j*+1_, where ∘ denotes the composition operator on relationships and 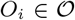 and 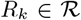 are object and relation type, respectively.

### Example 2.3.1.

MetaCyc contains multiple meta-paths conveying different semantics. For example, a metapath “ZCZ” represents the co-catalyst relationships on a compound (C) between two enzymatic reactions (Z), and “ZCTCZ” may indicate a meta-path that requires two enzymatic reactions (Z) transforming two compounds (C) within a pathway (T). Another important meta-path to consider is “CZC”, which implies “C + Z ⇒ C” transformation relationship.

### Problem Statement 1. Metabolic Pathway Prediction

*Given three inputs: i)-a heterogeneous information network* 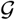, *ii)- a dataset* 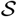, *and iii)- an optional set of meta-paths* 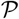, *the goal is to automatically resolve node embeddings such that leveraging the features will effectively improve pathway prediction for a hitherto unseeen instance* x*.

## 3 The pathway2vec Framework

The pathway2vec framework is composed of five modules: i)- node2vec ([16]), ii)- metapath2vec ([12]), iii)- metapath2vec++ ([12]), iv)- JUST ([18]), and v)- RUST (this work), where each module contains a random walk modeling and node representation step. A graphical representation of the pathway2vec framework is depicted in Fig. 2.

**C1. Random Walks.** In this step, a sequence of random walks over an input graph (whether heterogeneous or homogeneous) is generated based on the selected model. (see Section 3.1).
**C2. Learning Node Representation.** Resulting walks are fed into the Skip-Gram model to learn node embeddings ([25, 15, 16, 12]). An embedding is a low dimensional latent continuous feature for each node in 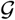, which encodes smooth decision boundaries between groups or communities within a graph. Details are provided in Section 3.2.

**Figure 2:**
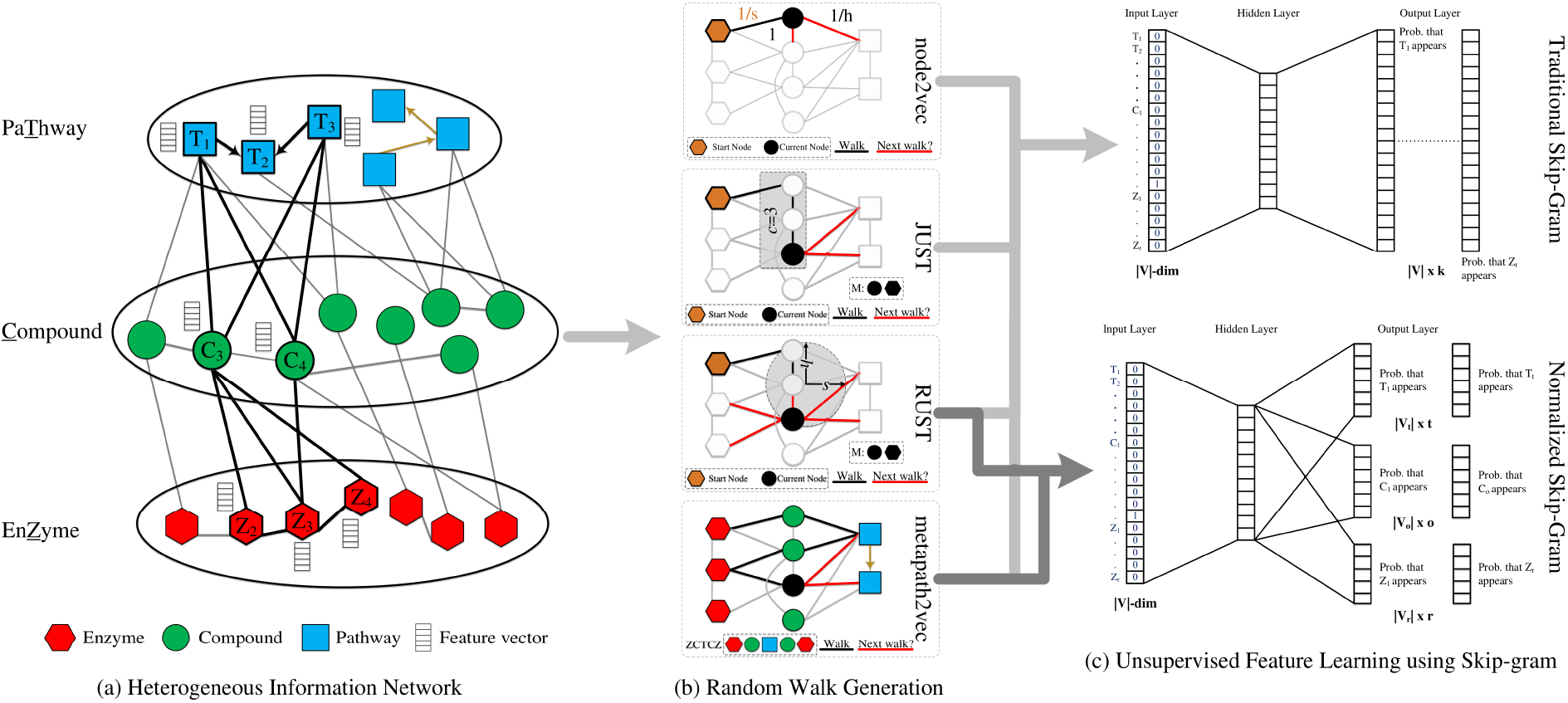
Graphical representation of pathway2vec framework. Main components: (a) a multi-layer heterogeneous information network composed from MetaCyc, showing meta-level interaction among compounds, enzymes, and pathways, (b) four random walks, and (c) two representational learning models: traditional Skip-Gram (top) and Skip-Gram by normalizing domain types (bottom). In the subfigure (a), the highlighted network neighbors of T_1_ (nitrifier denitrification) indicate this pathway interacts directly with T_2_ (nitrogen fixation I (ferredoxin)) and indirectly to T_3_ (nitrate reduction I (denitrification)) by second-order with relationships to several compounds, including nitric oxide (C_3_) and nitrite (C_4_) converted by enzymes represented by the EC numbers (Z_2_: EC 1.7.2.6, Z_3_: EC 1.7.2.1, and Z_4_: EC 1.7.2.5). The black colored nodes in subfigure (b) indicate the current position of the walkers and red links suggest the next possible nodes to sample while black links indicate route taken by a walker to reach the current node. node2vec is parameterized by local search *s* and in-out *h* hyperparameters. These two hyperparameters constitute a unit circle, i.e., *h*^2^ + *s*^2^ = 1, for RUST. *M* stores previously visited node types which is 2 and only applied for JUST and RUST. *c* is number of nodes of the same domain type as the current node which is 3 and is associated with JUST. For metapath2vec, a walker requires a prespecified scheme which is set to “ZCTCZ”. The normalized Skip-Gram in the subfigure (c) bottom is simply trained based on the domain type, in contrast to the traditional Skip-Gram model. More information related to both learning strategies is provided in Section 3.2. Zoom for readability.

### 3.1 Random Walks

To capture meaningful graph relationships, existing techniques such as DeepWalk ([28]), design simple but effective algorithms based on random walks for representational learning of features. However, DeepWalk does not address in-depth and in-breadth graph exploration. Therefore, node2vec ([16]) was developed to traverse local and global graph structures based on the principles of: i)- homophily ([26, 14]) where interconnected nodes form a community of correlated attributes and ii)- structural equivalence ([17]) where nodes having similar structural roles in a graph should be close to one another. node2vec simulates a second-order random walk, where the next node is sampled conditioned on the previous and the current node in a walk. For this, two hyper-parameters are adjusted, 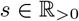 that extracts local information of a graph, and 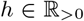 that enables local and global traversals by moving deep in a graph or walking within the vicinity of the current node. This method is illustrated in Fig. 2 (b) top.

First-order and second-order random walks were initially proposed for homogeneous graphs, but can be readily extended to heterogeneous information networks. Sun and colleagues ([31]) have observed that random walks can suffer from implicit bias due to initial node selection or the presence of a small set of dominant node types skewing results toward a subset of interconnected nodes. metapath2vec was developed ([12]), to resolve implicit bias in graph traversal to characterize semantic associations embodied between different types of nodes according to a certain path definition. This method is illustrated in Fig. 2 (b) bottom.

metapath2vec overcome the limitation of nove2vec by enabling to extract semantical representations over heterogeneous graph. However, the use of meta-paths requires either prior domain-specific knowledge to recover semantic associations of HIN according to a certain path definition. As a result, groups of vertices with the heterogeneous information network may not be visited or revisited multiple times. This limitation was partially addressed by leveraging multiple path schemes ([15]) to guide random walks based on a metapath length parameter. Hussein and colleagues developed the Jump and Stay (JUST) heterogeneous graph embedding method using random walks [18] as an alternative to meta-paths. JUST randomly selects the next node in a walk from either the same node type or from different node types using an exponential decay function and a tuning parameter based on on two history records: i)- *c* corresponding the number of nodes consecutively visited in the same domain as the current node and ii)- a queue *M* of size *m* storing the previously node types. This method is illustrated in Fig. 2 (b) second from top.

However, in order to balance the node distribution over multiple node types, JUST constrains the number of memorized domains *m* to be within the range of 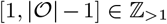. This can misrepresent graph structure in two ways: i)- explorations within domain because the last visited consecutive *c* nodes may enforce sampling from another domain, or ii) jumping deep towards nodes from other domains because *M* is constrained. To alleviate these problems we develop a novel random walk algorithm, RUST, adopting a unit-circle equation to sample node pairs that generalize previous representational learning methods, as illustrated in Fig. 2 (b) second from bottom. The two hyper-parameters *s* and *h* constitute a unit circle, i.e., *h*^2^ + *s*^2^ = 1, where *h* ∈ [0,1] indicates how much exploration is needed within a domain while *s* ∈ [0,1] defines the in-depth search towards other domains such that *s* > *h* encourages the walk to explore more domains and vice versa. Consequently, RUST blends both semantic associations and local/global structural information for generating walks without restricting domain size *m* in *M*.

To better illustrate the effect of *s* and *h* on RUST, consider an example in Fig. 3, where the walkers in JUST and RUST are currently stationed at *C*_3_ of compound type. While JUST enforces its walker to jump towards pathway domain, because of the combined effect of *c* that holds three consecutive nodes of compound type and *M* that is currently storing EC and compound types, RUST may prefer returning to *C*_2_ (no links exist to *C*_4_) than jumping to *T*_1_ or *T*_2_. This is because *s* < *h* entailing to explore more within the same domain as *C*_3_. If, however, *s* > *h* then RUST will perform in-depth search by selecting a node of type pathway. For formal definitions about the discussed random walks, see Supp. Section 1.

**Figure 3:**
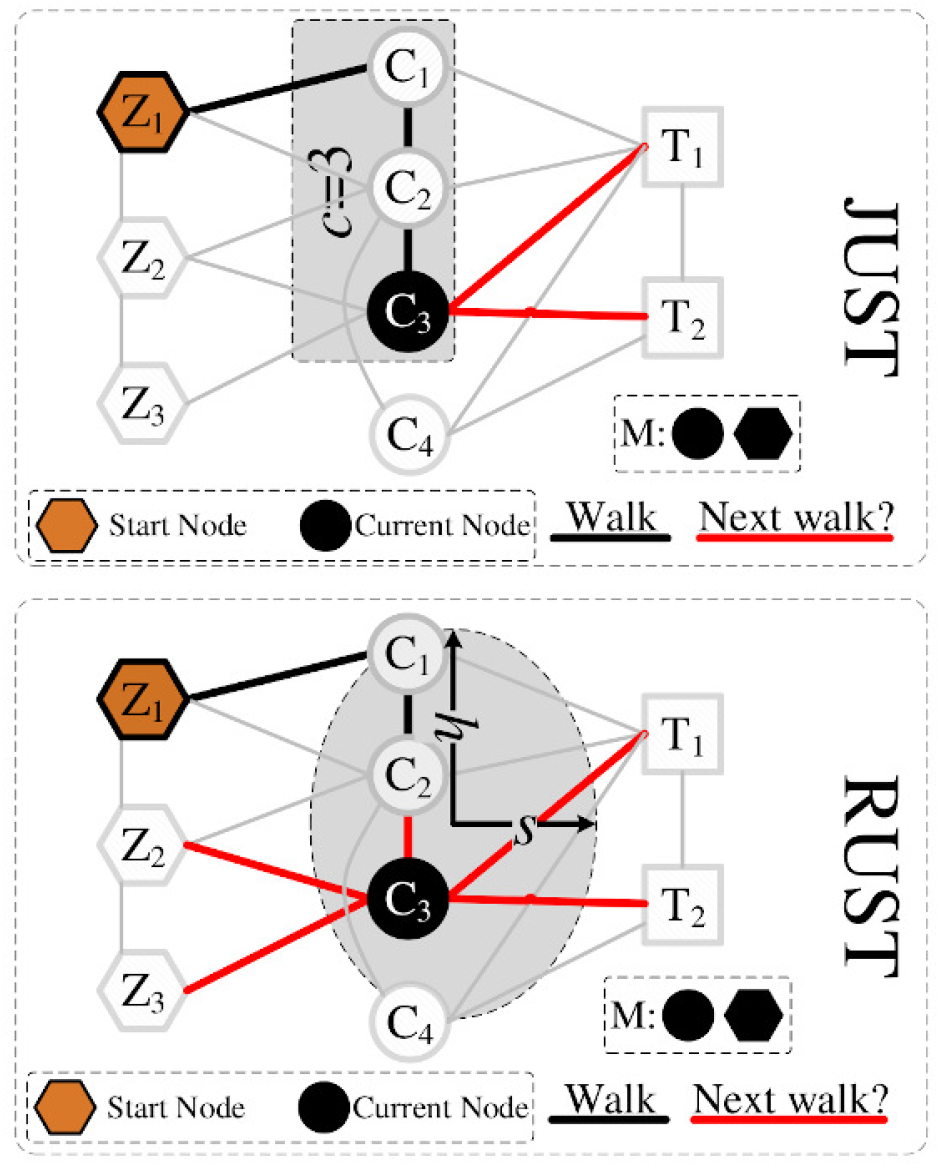
An illustrative example showing the selection of the next node for both JUST and RUST on HIN extracted from MetaCyc. The walker is currently stationed at *C*_3_ arriving from node *C*_2_ (indicated by black colored link), where *M* stores two previously visited node types and *c* (for JUST) holds 3 consecutive nodes that are of the same domain as *C*_3_. As can be seen JUST would prefer selecting the next node of type pathway while RUST may prefer returning to *C*_2_ than jumping to *T*_1_ or *T*_2_, as indicated by red edges, because *s* < *h* represented by an ellipsis glyph.

### 3.2 Learning Latent Embedding in Graph

Random walks 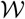 generated using node2vec, metapath2vec, JUST and RUST are fed into the Skip-Gram model to learn node embeddings ([25]). The Skip-Gram model exploits context information defined as a fixed number of nodes surrounding a target node. The model attempts to maximize co-occurrence probability among a pair of nodes identified within a given window of size *q* in 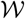 based on log-likelihood:

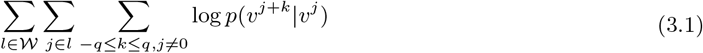

where *υ*^*j*–*c*^, …, *υ*^*j*+*c*^ are the context neighbor nodes of node *υ^j^* and *p*(*υ*^*j*+*i*^|*υ^j^*) defines the conditional probability of having context nodes given the node *υ^j^*. The *p*(*υ*^*j*+*k*^|*υ^j^*) is the commonly used softmax function, 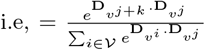where 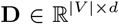 stores the embeddings of all nodes and **D**_*υ*_ is the *υ*-th row corresponding to the embedding vector for node *υ*. In practice, the vocabulary of nodes may be very large, which intensifies the computation of *p*(*υ*^*j*+*k*^|*υ^j^*). The Skip-Gram model uses negative sampling, which randomly selects a small set of nodes *N* that are not in the context to reduce computational complexity. This idea, represented in updated Eq. 3.1 is implemented in node2vec, metapath2vec, JUST, and RUST according to:

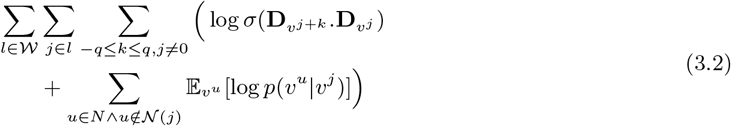

where 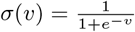 is the sigmoid function.

In addition to the equation above, Dong and colleagues proposed a normalized version of metapath2vec, called metapath2vec++, where the domain type of the context node is considered in calculating the probability *p*(*υ*^*j*+*k*^|*υ^j^*), resulting in the following objective formula:

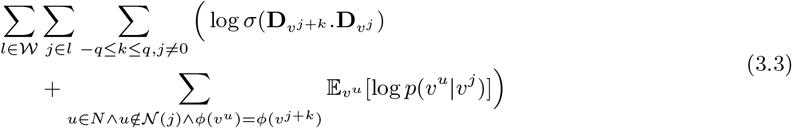

where *ϕ*(*υ^u^*) = *ϕ*(*υ*^*j*+*k*^) suggests that the negative nodes are of the same type as the context node *ϕ*(*υ*^*j*+*k*^). The above formula is also applied for RUST, and we refer it to RUST-norm. Through iterative update over all the context nodes, whether using Eq. 3.2 or Eq. 3.3, for each walk in 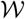, the learned features are expected to capture semantic and structural contents of a graph, thereby, can be made useful for pathway inference.

## 4 Predicting Pathways

For pathway inference, the learned EC embedding vectors are concatenated into each example *i* according to:

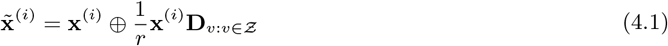

where ⊕ denotes the vector concatenation operation, 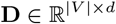 stores the embeddings of all nodes and 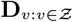 indicates feature vectors for *r* enzymatic reactions. By incorporating enzymatic reaction features into x^(*i*)^, the dimension size is extended to *r* + *d*, where *r* is the enzyme vector size while *d* corresponds to embeddings size. This modified version of x^(*i*)^ is denoted by 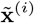, which then can be used by an appropriate machine learning algorithm, such as mlLGPR ([6]), to train and infer a set of metabolic pathways from enzymatic reactions.

## 5 Experimental Setup

In this section, we explain the experimental settings and outline materials used to evaluate the performance of pathway2vec modules that were written in Python v3 and trained using tensorflow v1.10 ([1]). Unless otherwise specified all tests were conducted on a Linux server using 10 cores of Intel Xeon CPU E5-2650.

### 5.1 Preprocessing MetaCyc

We constructed three hierarchical layers of HIN using MetaCyc v21 ([8]), according to: EC (bottom-layer), compound (mid-layer), and pathway (top-layer) as in Fig. 2(a). Relationships among these layers establish inter-interactions and betweenness interactions. Three inter-interactions were built: i)- ECs interactions that were collected based shared metabolites, e.g. if a compound is engaged in two ECs then the two ECs were considered connected; ii)- compounds interactions that were processed based on shared reactions, e.g. if any two compounds constituting substrate and product of an engaged enzymatic reaction they would be linked; and iii)- pathways interactions that were constructed based on shared metabolites, e.g. if any product in one pathway is being consumed by another then these two pathways were linked. With regard to betweenness interactions, we considered two forms: i)- EC-compound interaction if any enzyme (represented by an EC number) engages in any compound then nodes of both types were linked and ii)- compound-pathway interaction if any compound involves in any pathway then those nodes were considered related. After building multi-layer HIN, we apply different configurations, as summarized in Table 1, to explore the relationship between different graph types and the quality of generated walks and embeddings.

**Table 1:**
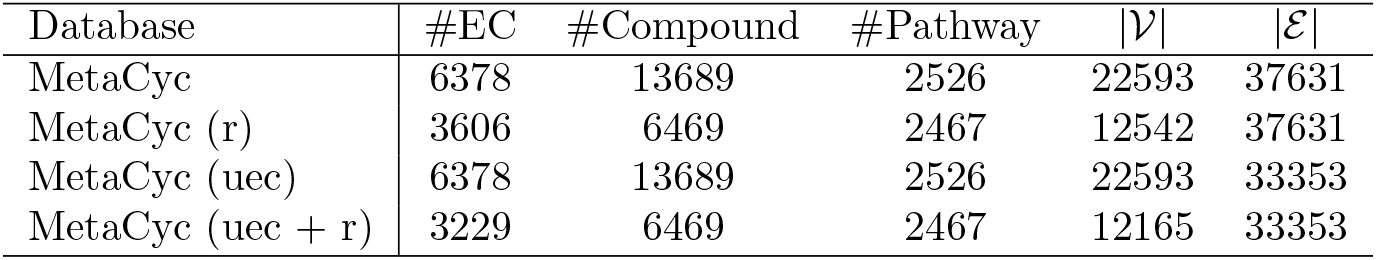
Different configurations of compound, enzyme (EC), and pathway objects extracted from the MetaCyc database: i)- full content (MetaCyc), ii)- reduced content based on trimming nodes below 2 links (MetaCyc r), iii)- links among enzymatic reactions are removed, following graph independence assumption (MetaCyc uec)), and iv)- combination of unconnected enzymatic reactions and trimmed nodes (MetaCyc uec + r).

| Database | #EC | #Compound | #Pathway | $\vert \mathcal{V}\vert $ | $\vert \mathcal{E}\vert $ |
| --- | --- | --- | --- | --- | --- |
| MetaCyc | 6378 | 13689 | 2526 | 22593 | 37631 |
| MetaCyc (r) | 3606 | 6469 | 2467 | 12542 | 37631 |
| MetaCyc (uec) | 6378 | 13689 | 2526 | 22593 | 33353 |
| MetaCyc (uec + r) | 3229 | 6469 | 2467 | 12165 | 33353 |

### 5.2 Parameter Settings

Parameterization for the other random walk methods can be found in ([16, 12, 18]). For training, we randomly initialized model parameters with a truncated Gaussian distribution, and set the learning rate to 0.01, the batch size to 100, and the number of epochs to 10. Unless otherwise indicated, for each module, the number of sampled path instances is *K* = 100, the walk length is *l* = 100, the embedding dimension size is *d* = 128, the neighborhood size is 5, the size of negative samples is 5, and the number of memorized domain *m* for JUST and RUST are 2 and 3, respectively. The explore and the in-out hyperparameters for node2vec and RUST are *h* = 0.7 (or *h* = 0.55) and *s* = 0.7 (or *s* = 0.84), respectively, using the uec configuration. For metapath2vec and metapath2vec++, we applied the meta-path scheme “ZCTCZ” to guide random walks. For brevity, we denote node2vec, metapath2vec, metapath2vec++, JUST, RUST, and RUST-norm as n2v, m2v, cm2v, jt, and rt, crt, respectively.

## 6 Experimental Results and Discussion

In this section, we first evaluate parameter sensitivity of RUST prior to benchmarking the four random walk algorithms, jointly with the two learning methods, based on node-clustering, embedding visualization, and pathway prediction.

### 6.1 Parameter Sensitivity of RUST

#### Experimental setup

In this section, the effect of different hyperparameter settings in RUST on the quality of learned nodes embeddings is described. Since the hyperparameter space involved in RUST, is infinite, exhaustive searches for optimal settings are prohibitive. Therefore, settings were sub-selected to determine RUST performance. Specifically, the effects of the dimensions *d* ∈ {30, 50, 80, 100, 128, 150}, the neighborhood size *q* ∈ {3, 5, 7, 9}, the memorized domains *m* ∈ {3, 5, 7}, and the two hyperparameters *s* and *h* (∈ {0.55, 0.71, 0.84}) were evaluated based on Normalized Mutual Information (NMI) scores, after 10 trials. The NMI produces scores between 0, indicating no mutual information exists, and 1, indicating node clusters (feature groups) are perfectly correlated based on class information: enzyme, compound, and pathway. Clustering was performed using the k-means algorithm ([5]) to group data based on the learned representations from RUST as described in ([12, 18]). Random walks 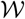 were generated using MetaCyc with uec option for RUST test parameters.

#### Experimental results

Fig. 4a indicates that RUST performance tends to saturate when the memorized domains are concentrated around *m* = 5 and *h* = 0.55, indicating a preference to explore more domain types. By fixing *m* = 3 and *h* = 0.55 the optimal results of NMI score w.r.t. the number of embedding dimensionality was found to be at 80 and 128 (Fig. 4b). Beyond this value RUST performance deteriorated. A similar trend was also observed when the context neighborhood size was increased beyond *q* > 5 (Fig. 4c). Based on these observations, the following settings *m* = 3, *h* = 0.55, *d* = 80 or *d* = 128, and *q* = 5 provide the most efficient and accurate clustering outcomes using MetaCyc with uec option. For comparative purposes, we set *d* = 128.

**Figure 4:**
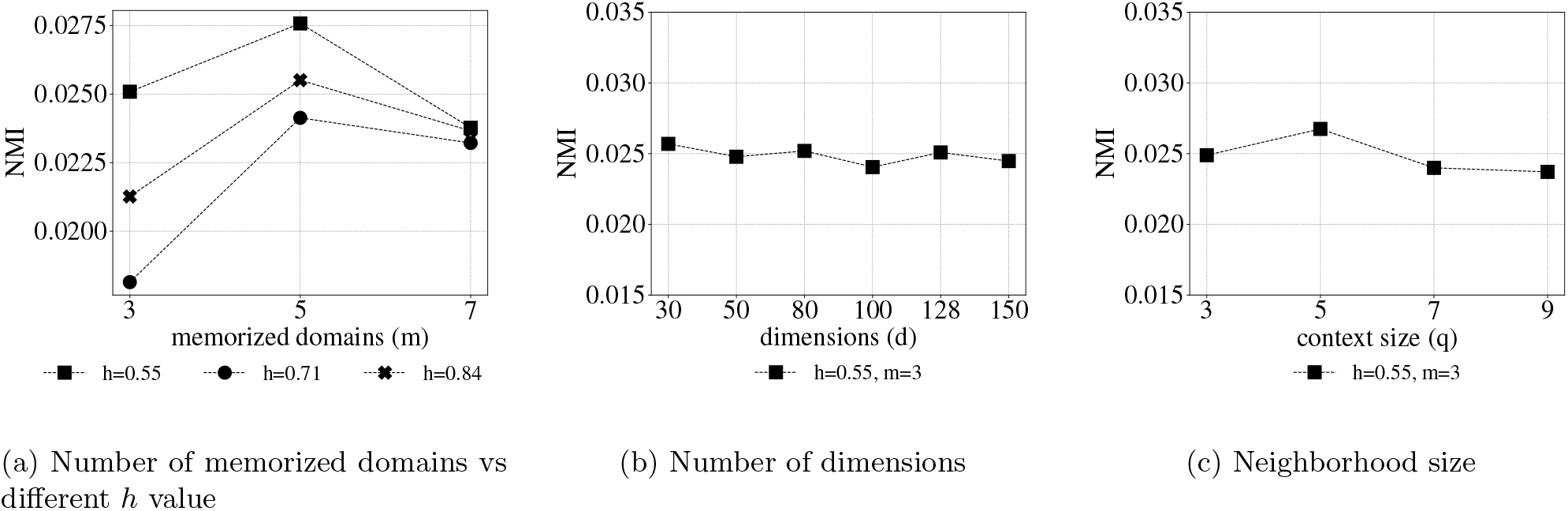
Parameter sensitivity of RUST based on NMI metric.

### 6.2 Node Clustering

#### Experimental setup

The performance of different random walk methods was tested in relation to node clustering using NMI after 10 trials and the hyperparameters described above on all MetaCyc graph types depicted in Table 1. Clustering was performed using the k-means algorithm to group homogeneous nodes based on the embeddings learned by each method.

#### Experimental results

Fig. 5 indicates node clustering results for node2vec, metapath2vec, JUST and RUST. node2vec, JUST and RUST exhibited similar performance across all configurations, indicating that these methods are less likely to extract semantic knowledge, characterizing node domains, from MetaCyc. However, RUST performed optimally better than node2vec and JUST in learning representations. In the case of metapath2vec, the random walk follows a predefined meta-path scheme, capturing the necessary relational knowledge for defining node types. For example, *nitrogenase* (EC-1.18.6.1), which reduces *nitrogen* gas into *ammonium*, is exclusively linked to the *nitrogen fixation I (ferredoxin)* pathway ([13]). Without a predefined relation, a walker may explore more local/global structure of 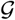, hence, become less efficient in exploiting relations between these two nodes. Among the four walks, only metapath2vec is able to accurately group those nodes, according to their classes. Despite the advantages of metapath2vec, it is biased to a scheme, as described in ([18]), which is explicitly observed for the case of “uec+r” (Fig. 5d). Under these conditions, both isolated nodes and links among ECs are discarded, resulting in a reduced number of nodes that are more easily traversed by a meta-path walker. metapath2vec++ exhibited trends similar to metapath2vec because they share the same walks. However, metapath2vec++ is trained using normalized Skip-Gram. Therefore, it is expected to achieve good NMI scores, yielding over 0.41 on uec+full content (in Supp. Section 2), which is also similar to RUST-norm NMI score (~ 0.38). This is interesting because RUST-norm employs RUST based walks but the embeddings are learned using normalized Skip-Gram.

**Figure 5:**
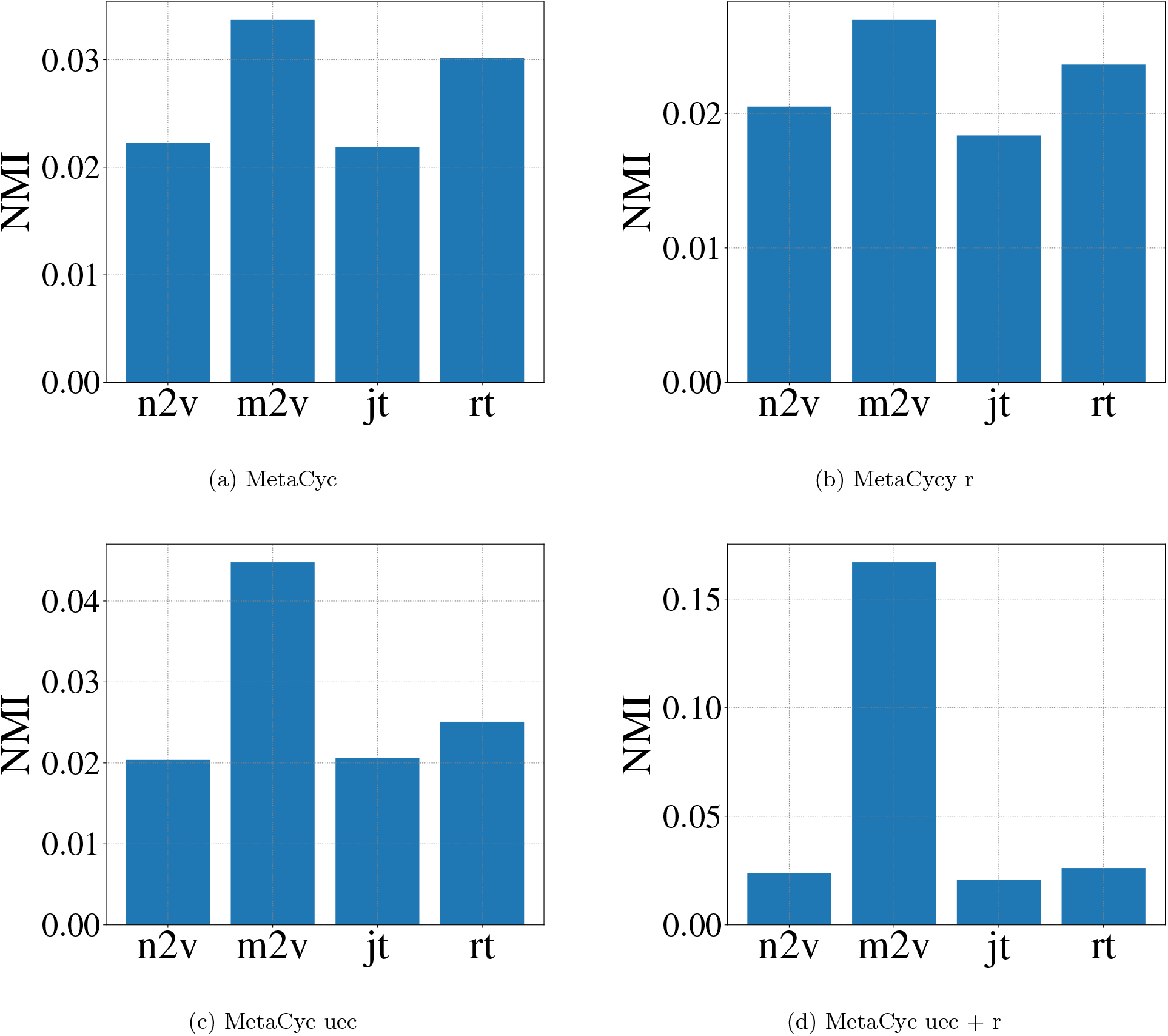
Node clustering results based on NMI metric using MetaCyc data.

Taken together, these results indicate that node2vec, JUST, and RUST based walks are effective for analyzing graph structure while metapath2vec can learn good embeddings. However, RUST strikes a balance between the two proprieties through proper adjustments of *m* and the two unit-circle hyperparameters. Regarding the MetaCyc type, we recommend “uec” because the associations among ECs are captured at the pathway level. The trimmed graph is contraindicated, because it eliminates many isolated, but important pathways and ECs.

### 6.3 Manifold Visualization

#### Experimental setup

In this section, learned high dimensional embeddings are visualized by projecting them onto a two-dimensional space using two case studies. The first case examines the quality of learned nodes embeddings according to the generated random walks an approach commonly sought in most graph learning embedding techniques ([16, 34]). We posit that a good representational learning method defines clear boundaries for nodes of the same type. For illustrative purposes, nodes corresponding to nitrogen metabolism were selected. The second case examines the limitations of meta-path based random walks, extending our discussions in Section 6.2. For illustrative purposes we focus on the pathway layer in Fig. 2a and consider representation of pathways having no enzymatic reactions. For visualization, we use UMAP, a.k.a. uniform manifold approximation and projection ([24]) using 1000 epochs with the remaining settings set to default values.

#### Experimental results

Fig. 6 visualizes 2D UMAP projections of the 128 dimension embeddings, trained under uec+full setting depicting 185 nodes related to nitrogen metabolism in MetaCyc. Each point denotes a node in HIN and each color indicates the node type. node2vec (Fig. 6a), JUST (Fig. 6c), and RUST (Fig. 6d) appear to be less than optimal in extracting walks that preserve three layer relational knowledge e.g. nodes belonging to different types form unclear boundaries and diffuse clusters. In the cases of metapath2vec (Fig. 6b), metapath2vec++ (Fig. 6f), and RUST-norm (Fig. 6f), nodes of the same color are more optimally portrayed. In the second use case 80 pathways were identified, having no enzymatic reactions, with their 109 pathway neighbors, as shown in Fig. 7a. From Fig. 7, we observe that, in contrast to node2vec, JUST, RUST, and RUST-norm, pathway nodes are skewed incorrectly in both metapath2vec and metapath2vec+—+ and (with lesser degree). This demonstrates the rigidness of meta-path based methods that follow a defined scheme that limits their capacity to exploit local structure in learning embeddings. Interestingly, RUST-norm, based on RUST walks, is the only method that combines structural and semantic information as indicated in Fig. 7g and Fig. 6f, respectively. Taken together, these results indicate that RUST based walks with training using Eq. 3.3 provide efficient embeddings, consistent with node clustering observations.

**Figure 6:**
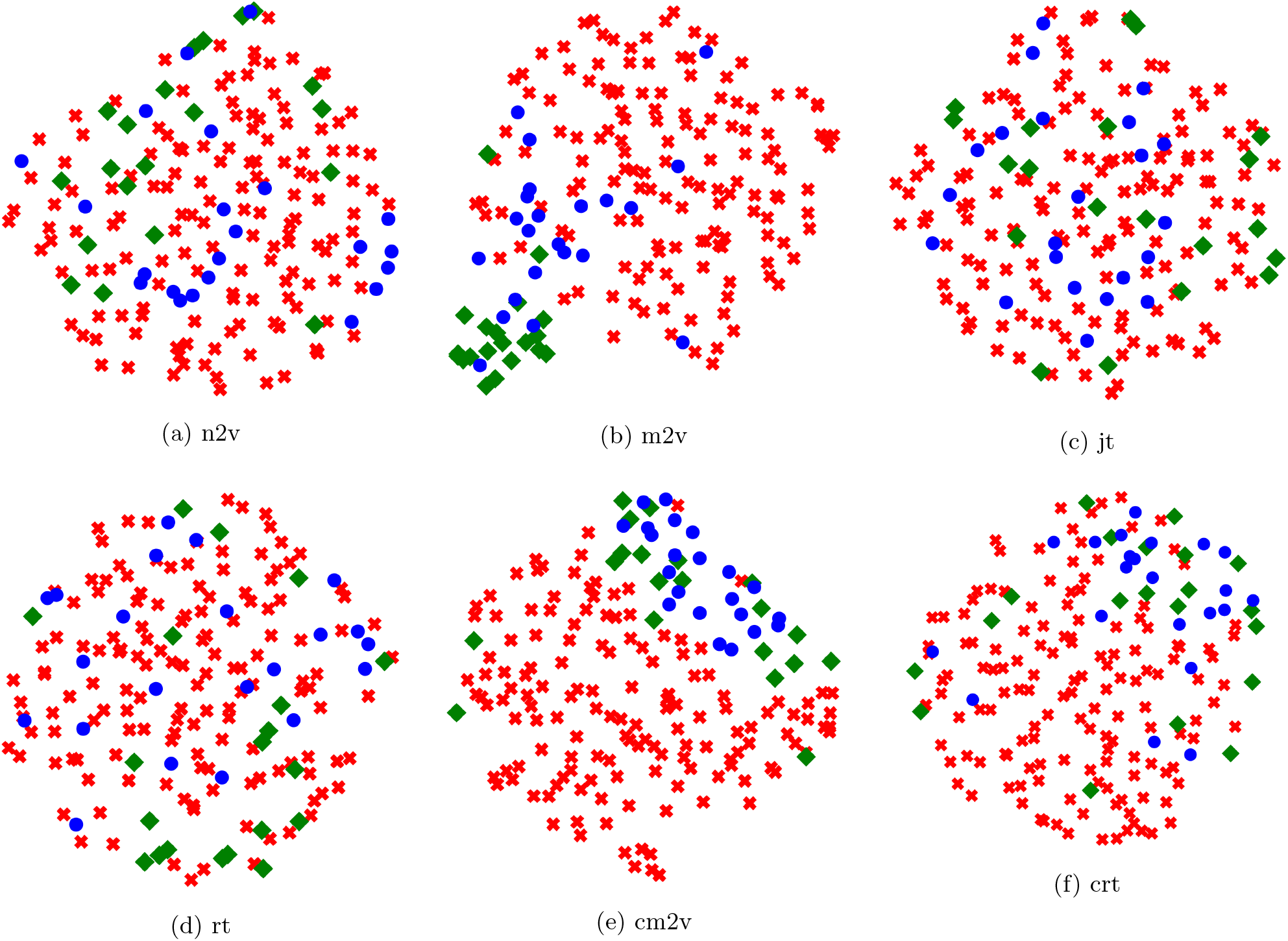
2D UMAP projections of the 128 dimension embeddings, trained under uec+full setting depicting 185 nodes related to nitrogen metabolism. Node color indicates the category of the node type, where red indicates enzymatic reactions, green indicates compounds, and blue is reserved for metabolic pathways.

**Figure 7:**
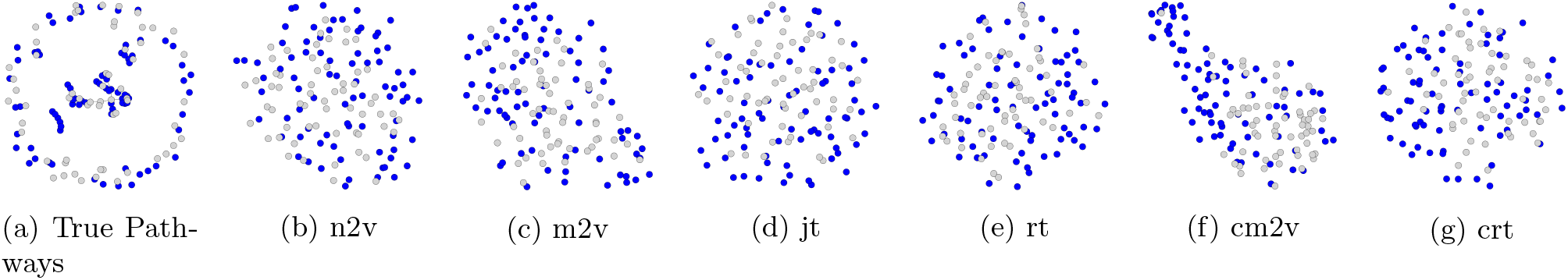
2D UMAP projections of 80 pathways that have no enzymatic reactions, indicated by the blue color, with 109 corresponding pathway neighbors, represented by the grey color.

### 6.4 Metabolic Pathway Prediction

#### Experimental setup

In this section, the effectiveness of the learned embeddings from pathway2vec modules is determined across different pathway inference methods including MinPath v1.2 ([35]), PathoLogic v21 ([21]), and mlLGPR-elastic net (EN), [6]. In contrast to previous multi-label classification methods ([28, 16, 18]), where the goal is to predict the most probable label set for nodes, we leverage the learned vectors and the multi-label dataset, according to Eq. 4.1. Pathway prediction with mlLGPR-EN used the default hyperparameter settings, after concatenating features from each learning method, to train on BioCyc (v20.5 tier (T) T2 & T3) ([9]) consisting of 9255 Pathway/Genome Databases (PGDBs) with 1463 distinct pathway labels (see Supp. Section 4). Results are reported on T1 golden datasets including EcoCyc, HumanCyc, AraCyc, YeastCyc, LeishCyc, and TrypanoCyc. Four evaluation metrics are used to report performance scores after 3 repeated trials: *Hamming loss, micro precision, micro recall*, and *micro F1 score*.

#### Experimental results

Table 2 shows micro F1 scores for each pathway predictor. Numbers in boldface represent the best performance score in each column while the underlined text indicates the best performance among the embedding methods. From the results, it is obvious that all variation of embedding methods performs consistently better than MinPath across the T1 golden datasets. With the excpetion of EcoCyc the performance of embeddings resulted in less optimal micro F1 scores than PathoLogic or mlLGPR. In the case of mlLGPR, embeddings were trained on less than 1470 pathways, potentially obscuring the actual benefits of the learned features. Taken together, different pathway2vec modules performed similar to one another indicating that embeddings are potential alternatives to the pathway and reaction evidence features used in ([6]). Full results are provided in Supp. Section 5.

**Table 2.**
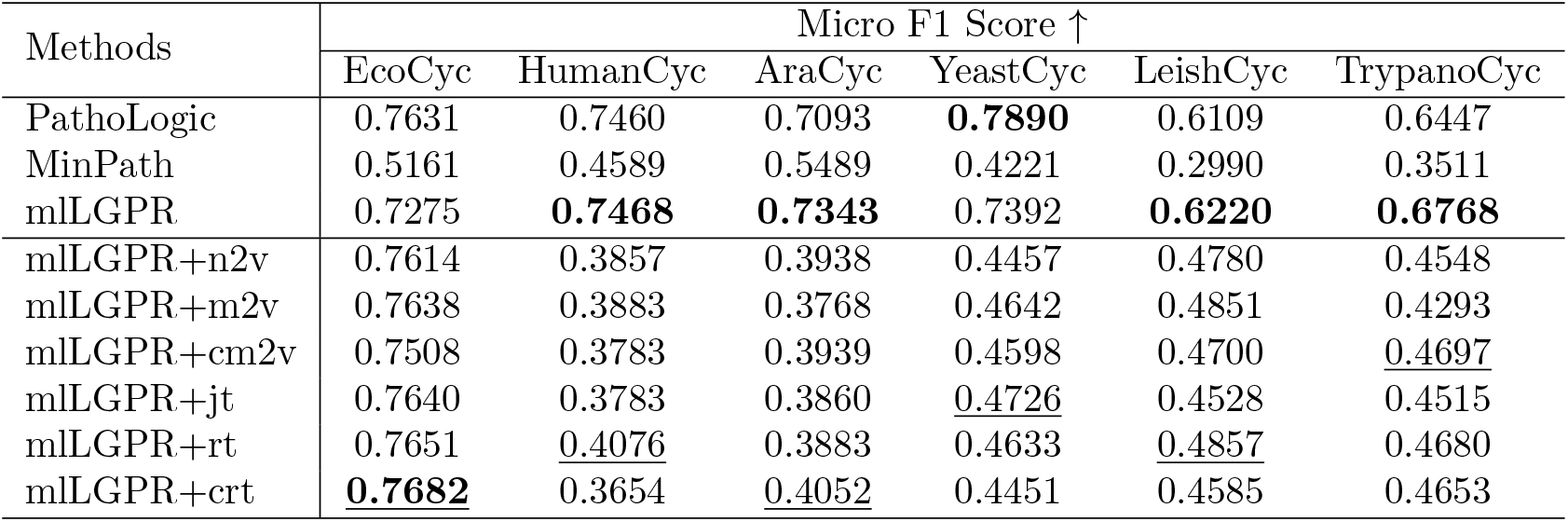
Micro F1 scores of each comparing algorithm on 6 benchmark datasets.

## 7 Conclusion

We have developed the pathway2vec package for learning features relevant to metabolic pathway prediction from genomic sequence information. The software package consists of six representational learning modules used to automatically generate features for pathway inference. Metabolic feature representations were decomposed into three interacting layers: compounds, enzymes and pathways, where each layer consists of associated nodes. A Skip-Gram model was applied to extract embeddings for each node encoding smooth decision boundaries between groups of nodes in a graph resulting in a multi-layer heterogeneous information network for metabolic interactions within and between layers. Three extensive empirical studies were conducted to benchmark pathway2vec, indicating that the representational learning approach is a promising adjunct or alternative to features engineering based on manual curation. At the same time, we introduced RUST, a novel and flexible random walk method that uses unit-circle and domain size hyperparameters to exploit local/global structure while absorbing semantic information from both homogeneous and heterogeneous graphs. Looking forward, we intend to leverage embeddings and graph structure on more complex community level metabolic pathway prediction problems. Because random walk based methods depend on many hyperparameters (e.g. the length of a random walk) that must be tuned, and many walks that must be generated we are exploring alternative graph convolutional neural networks to reduce computational complexity. Such methods aggregate feature information based on node co-occurrences patterns automatically without dependence on hyperparameter settings ([2, 10, 27]).

## Acknowledgments

We would like to thank Connor Morgan-Lang, Julia Glinos, Kishori Konwar and Aria Hahn for lucid discussions on the function of the pathway2vec framework, Ryan MacLaughlin for his participation in preliminary performance evaluations and all members of the Hallam Lab for helpful comments along the way.

## Funding

This work was performed under the auspices Genome Canada, Genome British Columbia, the Natural Science and Engineering Research Council (NSERC) of Canada, and Compute/Calcul Canada). ARB was supported by a UBC four-year doctoral fellowship (4YF) administered through the UBC Graduate Program in Bioinformatics.

## Conflict of Interest

none declared.

## Notes

#### Summary of Updates

The content of the manuscript is updated.

https://zenodo.org/record/3711103#.Xnlj-XVKjeR

## References

[1] Martin Abadi, Paul Barham, Jianmin Chen, Zhifeng Chen, Andy Davis, Jeffrey Dean, Matthieu Devin, Sanjay Ghemawat, Geoffrey Irving, Michael Isard, et al. Tensorflow: A system for large-scale machine learning. In 12th {USENIX} Symposium on Operating Systems Design and Implementation ({OSDI} 16), pp. 265–283, 2016.

[2] Sami Abu-El-Haija, Bryan Perozzi, Rami Al-Rfou, and Alexander A Alemi. Watch your step: Learning node embeddings via graph attention. In Advances in Neural Information Processing Systems, pp. 9180–9190, 2018.

[3] Sahar Abubucker, Nicola Segata, Johannes Goll, Alyxandria M Schubert, Jacques Izard, Brandi L Cantarel, Beltran Rodriguez-Mueller, Jeremy Zucker, Mathangi Thiagarajan, Bernard Henrissat, et al. Metabolic reconstruction for metagenomic data and its application to the human microbiome. PLoS Comput Biol, 8(6):e1002358, 2012.

[4] Wilhelm J Ansorge. Next-generation dna sequencing techniques. New biotechnology, 25(4):195–203, 2009.

[5] David Arthur and Sergei Vassilvitskii. k-means++: The advantages of careful seeding. In Proceedings of the eighteenth annual ACM-SIAM symposium on Discrete algorithms, pp. 1027–1035. Society for Industrial and Applied Mathematics, 2007.

[6] Abdur Rahman MA Basher, Ryan J McLaughlin, and Steven J Hallam. Metabolic pathway inference using multi-label classification with rich pathway features. bioRxiv, 2020.

[7] Pablo Carbonell, Jerry Wong, Neil Swainston, Eriko Takano, Nicholas J Turner, Nigel S Scrutton, Douglas B Kell, Rainer Breitling, and Jean-Loup Faulon. Selenzyme: Enzyme selection tool for pathway design. Bioinformatics, 34(12):2153–2154, 2018.

[8] Ron Caspi, Richard Billington, Luciana Ferrer, Hartmut Foerster, Carol A. Fulcher, Ingrid M. Keseler, Anamika Kothari, Markus Krummenacker, Mario Latendresse, Lukas A. Mueller, Quang Ong, Suzanne Paley, Pallavi Subhraveti, Daniel S. Weaver, and Peter D. Karp. The metacyc database of metabolic pathways and enzymes and the biocyc collection of pathway/genome databases. Nucleic Acids Research, 44(D1):D471–D480, 2016.

[9] Ron Caspi, Richard Billington, Hartmut Foerster, Carol A Fulcher, Ingrid Keseler, Anamika Kothari, Markus Krummenacker, Mario Latendresse, Lukas A Mueller, Quang Ong, et al. Biocyc: Online resource for genome and metabolic pathway analysis. The FASEB Journal, 30(1 Supplement):lb192–lb192, 2016.

[10] Taco Cohen, Maurice Weiler, Berkay Kicanaoglu, and Max Welling. Gauge equivariant convolutional networks and the icosahedral cnn. In International Conference on Machine Learning, pp. 1321–1330, 2019.

[11] Joseph M Dale, Liviu Popescu, and Peter D Karp. Machine learning methods for metabolic pathway prediction. BMC bioinformatics, 11(1):1, 2010.

[12] Yuxiao Dong, Nitesh V Chawla, and Ananthram Swami. metapath2vec: Scalable representation learning for heterogeneous networks. In Proceedings of the 23rd ACM SIGKDD International Conference on Knowledge Discovery and Data Mining, pp. 135–144. ACM, 2017.

[13] Robert R Eady. Structure- function relationships of alternative nitrogenases. Chemical reviews, 96(7):3013–3030, 1996.

[14] Santo Fortunato. Community detection in graphs. Physics reports, 486(3-5):75–174, 2010.

[15] Tao-yang Fu, Wang-Chien Lee, and Zhen Lei. Hin2vec: Explore meta-paths in heterogeneous information networks for representation learning. In Proceedings of the 2017 ACM on Conference on Information and Knowledge Management, pp. 1797–1806. ACM, 2017.

[16] Aditya Grover and Jure Leskovec. node2vec: Scalable feature learning for networks. In Proceedings of the 22nd ACM SIGKDD international conference on Knowledge discovery and data mining, pp. 855–864. ACM, 2016.

[17] Keith Henderson, Brian Gallagher, Tina Eliassi-Rad, Hanghang Tong, Sugato Basu, Leman Akoglu, Danai Koutra, Christos Faloutsos, and Lei Li. Rolx: structural role extraction & mining in large graphs. In Proceedings of the 18th ACM SIGKDD international conference on Knowledge discovery and data mining, pp. 1231–1239. ACM, 2012.

[18] Rana Hussein, Dingqi Yang, and Philippe Cudre-Mauroux. Are meta-paths necessary?: Revisiting heterogeneous graph embeddings. In Proceedings of the 27th ACM International Conference on Information and Knowledge Management, pp. 437–446. ACM, 2018.

[19] Dazhi Jiao, Yuzhen Ye, and Haixu Tang. Probabilistic inference of biochemical reactions in microbial communities from metagenomic sequences. PLoS Comput Biol, 9(3):e1002981, 2013.

[20] Minoru Kanehisa, Miho Furumichi, Mao Tanabe, Yoko Sato, and Kanae Morishima. Kegg: new perspectives on genomes, pathways, diseases and drugs. Nucleic Acids Research, 45(D1):D353–D361, 2017.

[21] Peter D Karp, Mario Latendresse, Suzanne M Paley, Markus Krummenacker, Quang D Ong, Richard Billington, Anamika Kothari, Daniel Weaver, Thomas Lee, Pallavi Subhraveti, et al. Pathway tools version 19.0 update: software for pathway/genome informatics and systems biology. Briefings in bioinformatics, 17(5):877–890, 2016.

[22] Peter D Karp, Wai Kit Ong, Suzanne Paley, Richard Billington, Ron Caspi, Carol Fulcher, Anamika Kothari, Markus Krummenacker, Mario Latendresse, Peter E Midford, et al. The ecocyc database. EcoSal Plus, 8(1), 2018.

[23] Christopher E Lawson, William R Harcombe, Roland Hatzenpichler, Stephen R Lindemann, Frank E Löffler, Michelle A O’Malley, Hector Garcia Martin, Brian F Pfleger, Lutgarde Raskin, Ophelia S Venturelli, et al. Common principles and best practices for engineering microbiomes. Nature Reviews Microbiology, pp. 1–17, 2019.

[24] Leland McInnes, John Healy, Nathaniel Saul, and Lukas Grossberger. Umap: Uniform manifold approximation and projection. The Journal of Open Source Software, 3(29):861, 2018.

[25] Tomas Mikolov, Ilya Sutskever, Kai Chen, Greg S Corrado, and Jeff Dean. Distributed representations of words and phrases and their compositionality. In Advances in neural information processing systems, pp. 3111–3119, 2013.

[26] Mark EJ Newman. Modularity and community structure in networks. Proceedings of the national academy of sciences, 103(23):8577–8582, 2006.

[27] Hongbin Pei, Bingzhe Wei, Kevin Chen-Chuan Chang, Yu Lei, and Bo Yang. Geom-gcn: Geometric graph convolutional networks. arXiv preprint arXiv:2002.05287, 2020.

[28] Bryan Perozzi, Rami Al-Rfou, and Steven Skiena. Deepwalk: Online learning of social representations. In Proceedings of the 20th ACM SIGKDD international conference on Knowledge discovery and data mining, pp. 701–710. ACM, 2014.

[29] Mahdi Shafiei, Katherine A Dunn, Hugh Chipman, Hong Gu, and Joseph P Bielawski. Biomenet: A bayesian model for inference of metabolic divergence among microbial communities. PLoS Comput Biol, 10(11):e1003918, 2014.

[30] Chuan Shi, Yitong Li, Jiawei Zhang, Yizhou Sun, and S Yu Philip. A survey of heterogeneous information network analysis. IEEE Transactions on Knowledge and Data Engineering, 29(1):17–37, 2017.

[31] Yizhou Sun, Jiawei Han, Xifeng Yan, Philip S Yu, and Tianyi Wu. Pathsim: Meta path-based top-k similarity search in heterogeneous information networks. Proceedings of the VLDB Endowment, 4(11), 2011.

[32] Yasuo Tabei, Yoshihiro Yamanishi, and Masaaki Kotera. Simultaneous prediction of enzyme orthologs from chemical transformation patterns for de novo metabolic pathway reconstruction. Bioinformatics, 32(12):i278–i287, 2016.

[33] David Toubiana, Rami Puzis, Lingling Wen, Noga Sikron, Assylay Kurmanbayeva, Aigerim Soltabayeva, Maria del Mar Rubio Wilhelmi, Nir Sade, Aaron Fait, Moshe Sagi, et al. Combined network analysis and machine learning allows the prediction of metabolic pathways from tomato metabolomics data. Communications Biology, 2(1):214, 2019.

[34] Daixin Wang, Peng Cui, and Wenwu Zhu. Structural deep network embedding. In Proceedings of the 22nd ACM SIGKDD international conference on Knowledge discovery and data mining, pp. 1225–1234. ACM, 2016.

[35] Yuzhen Ye and Thomas G Doak. A parsimony approach to biological pathway reconstruction/inference for genomes and metagenomes. PLoS Comput Biol, 5(8):e1000465, 2009.

