## Supplementary Material for "Leveraging Heterogeneous Network Embedding for Metabolic Pathway Prediction"

March 23, 2020

#### 1 Definitions

The following is the definition of a first-order random walk as implemented in DeepWalk [7], extending a nodes immediate neighbors to include nodes that are locally connected.

**Definition 1.1. Random Walk** [7]. A random walk $W$ of length $l$, rooted at node $v$, is a stochastic process with random variables $v^1, v^2, ..., v^l, v^{l+1}$ such that $v^{j+1}$ is a vertex sampled at random from the neighbors of vertex $v^j$ for all $1 \leq j \leq l$ according to the following distribution:

$$p(v^{j+1}|v^j) = \begin{cases} \frac{\alpha \pi_{j,j+1}}{Q} & \text{if } (v^j, v^{j+1}) \in \mathcal{E} \\ 0 & \text{otherwise} \end{cases} \tag{1.1}$$

where $\pi_{j,j+1} \in \mathbb{R}^{|\mathcal{V}| \times |\mathcal{V}|}$ is an unnormalized transition probability, indicating the probability of a random walker visiting a node $v^{j+1}$ conditioned on the current node being $v^j$, $Q$ is a normalizing term, and $\alpha \in [0,1]$ is a prior probability.

The above definition does not address in-depth and in-breadth graph exploration. Therefore, node2vec [4] was proposed where the process of node2vec random walks can be defined by manipulating $\alpha$ in Def. 1.1 according to:

$$\alpha_{s,h}(j-1, j+1) = \begin{cases} \frac{1}{s} & \text{if } \beta_{j-1,j+1} = 0 \\ 1 & \text{if } \beta_{j-1,j+1} = 1 \\ \frac{1}{h} & \text{if } \beta_{j-1,j+1} = 2 \end{cases} \tag{1.2}$$

where $\beta_{j-1,j+1} \in \{0, 1, 2\}$ denotes the distance between the previously visited node $v^{j-1}$ and the next neighbor node $v^{j+1}$.

First-order and second-order random walks were initially proposed for homogeneous graphs, but can be readily extended to heterogeneous information networks. This lead to developing metapath2vec [3] where meta-path based walks can be defined as:

**Definition 1.2. Meta-Path based Random Walk** [3]. Given a meta-path $P \in \mathcal{P}$ and $\mathcal{G}$, a meta-path based random walk $W$ of length $l$, rooted at node $v \in O$ as dictated by $P$, is a stochastic process with random variables $v_1^1, v_2^2, ..., v_k^l, v_{k+1}^{l+1}$ such that $v_{k+1}^{j+1}$ is a vertex of type $O_{k+1}$ sampled randomly from the neighbors of vertex $v_k^j \in O_k$ for all $1 \leq j \leq l$ according to the following distribution:

$$p(v^{j+1}|v_k^j, P) = \begin{cases} \frac{1}{|\mathcal{N}_{k+1}(v_k^j)|} & \text{if } (v_k^j, v^{j+1}) \in \mathcal{E}, \phi(v^{j+1}) = k+1 \\ 0 & \text{if } (v_k^j, v^{j+1}) \in \mathcal{E}, \phi(v^{j+1}) \neq k+1 \\ 0 & \text{if } (v_k^j, v^{j+1}) \notin \mathcal{E} \end{cases} \tag{1.3}$$

where $v_k^j \in \mathcal{O}_k$ and $\mathcal{N}_{k+1}(v_k^j)$ denotes the neighbors of $v_k^j$ that are of type $\mathcal{O}_{k+1}$ as pre-specified by the meta-path scheme $P$.

metapath2vec overcome the limitation of nove2vec by enabling to extract semantical representations over heterogeneous graph. However, the use of meta-paths requires either prior domain-specific knowledge to recover semantic associations of HIN according to a certain path definition. Hussein and colleagues developed the Jump and Stay (JUST) heterogeneous graph embedding method using random walks [5] as an alternative to meta-paths. JUST randomly selects the next node in a walk from either the same node type or from different node types using an exponential decay function and a tuning parameter.

**Definition 1.3. Jump and Stay based Random Walk (JUST)** [5]. Given a set of domain types $\mathcal{O}$, a graph $\mathcal{G}$, a queue $M$ of size $m$, and an initial stay probability $\alpha \in [0, 1]$, a JUST based random walk $W$ of length $l$, rooted at node $v \in O$, is a stochastic process with random variables $v^1, v^2, ..., v^l, v^{l+1}$ such that $v^{j+1}$ is selected according to the two following consecutive steps:

1. Predict the *stay probability* $p^{stay}$ as:

$$p^{\mathbf{st}}(v^j) = \begin{cases} 0 & \text{if } S(v^j) = \varnothing \\ 1 & \text{if } J(v^j) = \varnothing \\ \alpha^c & \text{if otherwise} \end{cases} \tag{1.4}$$

   where $c$ is the number of nodes consecutively visited in the same domain as $v^j$ and the remaining terms $S(v^j)$ and $J(v^j)$ are:

$$S(v^j) = \{v^{j+1} | (v^j, v^{j+1}) \in \mathcal{E} \wedge \phi(v^j) = \phi(v^{j+1})\}$$
$$J(v^j) = \{v^{j+1} | (v^j, v^{j+1}) \in \mathcal{E} \wedge \phi(v^j) \neq \phi(v^{j+1})\}$$

2. Sample $v^{j+1}$ either: i)- from the same domain as $v^j$ or ii)- apply the equation below, iff $p^{\mathbf{st}}(v^j) = 0$ or $p^{\mathbf{st}}(v^j) = 1 - \alpha^c$:

$$H(v^j) = \begin{cases} \{k | k \in \mathcal{O} \wedge k \notin T, J(v^j) \neq \varnothing\} & \text{if not empty} \\ \{k | k \in \mathcal{O}, k \neq \phi(v^j), J(v^j) \neq \varnothing\} & \text{if otherwise} \end{cases} \tag{1.5}$$

   where $M$ is a queue of size $m$ that stores $m$ previously visited types.

If the set $S(v^j)$ is empty, meaning no edges exist between $v^j$ and any nodes in $\mathcal{V}$ that share the same domain type as $v^j$, then a node is sampled from different types based on Eq. 1.5, which suggests to select randomly any domains not included in $M$. If, however, the latter condition is not satisfied then simply choose one domain that is different than the current node type. If $J(v^j)$ is empty, i.e., no heterogeneous edges connected to $v^j$, then the random walker is forced to stay in the same domain. Finally, if both homogeneous and heterogeneous edges are connected to $v^j$ then the walker may choose to either stay with $\alpha^c$ or jump with $1 - \alpha^c$, where $\alpha$ value decays exponentially by $c$ that stores the number of nodes sequentially visited in the same type of $v^j$. This can misrepresent graph structure in two ways: i)- explorations within domain because the last visited consecutive $c$ nodes may enforce sampling from another domain, or ii) jumping deep towards nodes from other domains because $M$ is constrained. To alleviate these problems we develop a novel random walk algorithm, RUST, adopting a unit-circle equation to sample node pairs that generalize previous representational learning methods.

**Definition 1.4. Unit-Circle based Jump and Stay Random Walk (RUST).** Given a set of domain types $\mathcal{O}$, a graph $\mathcal{G}$, a queue $M$ of size $m$, and two hyper-parameters $s$ and $h$, a RUST based random walk $W$ of length $l$, rooted at node $v \in O$, is a stochastic process with random variables $v^1, v^2, ..., v^l, v^{l+1}$ such that $v^{j+1}$ is chosen in two steps:

1. Estimate domain types transition probabilities given $v^j$:

$$\pi^{\mathbf{dom}}_{j,j+1} = \begin{cases} \frac{h.\beta_j \pi_{j-1,j}}{Q} & \text{if } S(v^j) = \varnothing \\ \frac{s.\beta_j \pi_{j-1,j}}{Q} & \text{if } J(v^j) = \varnothing \end{cases} \tag{1.6}$$

   where $Q$ is a normalizing term, $\beta_j \in (0, 1]$ is a domain weight hyperparameter of $v^j$ added to give more weights, if necessary, to some domains, and $\pi_{j-1,j}$ is an unnormalized transition probability from previous node $v^{j-1}$ to the current node $v^j$. The remaining terms:

$$S(v^j) = \{v^{j+1} | (v^j, v^{j+1}) \in \mathcal{E} \wedge \phi(v^j) = \phi(v^{j+1})\}$$
$$J(v^j) = \{v^{j+1} | (v^j, v^{j+1}) \in \mathcal{E} \wedge \phi(v^j) \neq \phi(v^{j+1})\}$$

2. Sample a domain type $k$ at random from $\pi^{\mathbf{dom}}_{j,j+1}$ according to:

$$H(v^j) = \{k | k \in \mathcal{O}, \alpha_k.\pi^k_{j,j+1}\} \tag{1.7}$$

   where $\alpha_k = 1/e^{c_k}$ and $c_k$ is the number of nodes with type $k$ that is stored in $M$. Finally, select randomly a node $v^{j+1}$ based on $H(v^j)$.

The two hyper-parameters $s$ and $h$ constitute a unit circle, i.e., $h^2 + s^2 = 1$, where $h \in [0, 1]$ indicates how much exploration is needed within a domain while $s \in [0, 1]$ defines the in-depth search towards other domains such that $s > h$ encourages the walk to explore more domains and vice versa. Consequently, RUST blends both semantic associations and local/global structural information for generating walks without restricting the number of memorized domains $m$ while the $\alpha_k$ serves as a function of node size having the type $k$ as stored in $M$. Algorithm 1 presents the pseudocode of RUST based random walk.

**Inputs** : A graph $\mathcal{G} = (\mathcal{V}, \mathcal{E})$, a prior probability $\alpha$, number of memorized domains $m$, explore hyperparameter $h$, in-out hyperparameter $s$, walk length $l$, number of random walks per node $K$
**Outputs:** A set of walks $\mathcal{W}$
**Process :**

Initialize a type transition probability $\pi^p$ over all nodes;
**for** $v \in \mathcal{V}$ **do**
    **for** $i \leftarrow 1$ **to** $K$ **do**
        `walk`$=[v]$;
        $M \leftarrow \varnothing$;
        **for** $j \leftarrow 1$ **to** $l - 1$ **do**
            $\pi^{\text{dom}}_{j,j+1} \leftarrow$ by applying Eq. 1.6;
            $H(v^j) \leftarrow$ by applying Eq. 1.7;
            **if** $|M| = m$ **then**
                Update $\alpha_k = 1/e^{c_k}$;
            Sample a $v^j$ from $H(v^j)$;
            `walk.append(`$v^j$`)`
    Add `walk` to $\mathcal{W}$;
Return $\mathcal{W}$;

**Algorithm 1:** RUST based Random Walk

### 2 Node Clustering

Fig. 1 shows the clustering results based on NMI score using metapath2vec++ (cm2v) and RUST-norm (crt) on four MetaCyc settings. Since metapath2vec++ is trained using normalized Skip-Gram, it is expected to achieve good NMI scores, yielding over 0.41 on uec+full content, which is also similar to RUST-norm NMI score ($\sim 0.38$). This is interesting because RUST-norm employs RUST based walks but the embeddings are learned using normalized Skip-Gram.

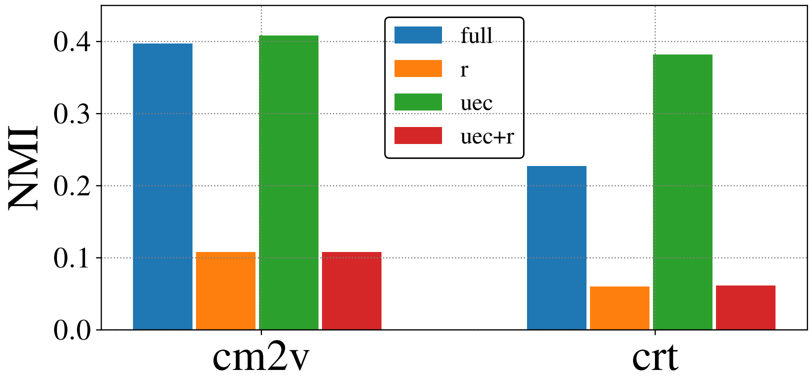


Figure 1: Node clustering results of metapath2vec++ (cm2v) and RUST-norm (crt) based on NMI metric using MetaCyc data.

### 3 Manifold visualization

Fig. 2 depicts ECs, compounds, and pathways, and their interactions, as extracted from MetaCyc. Also, we illustrate in Fig 4 the visualization of embeddings of 3000 randomly subsampled nodes, represented in Fig. 3, learned using node2vec, metapath2vec, metapath2vec++, JUST, RUST, and RUST-norm.

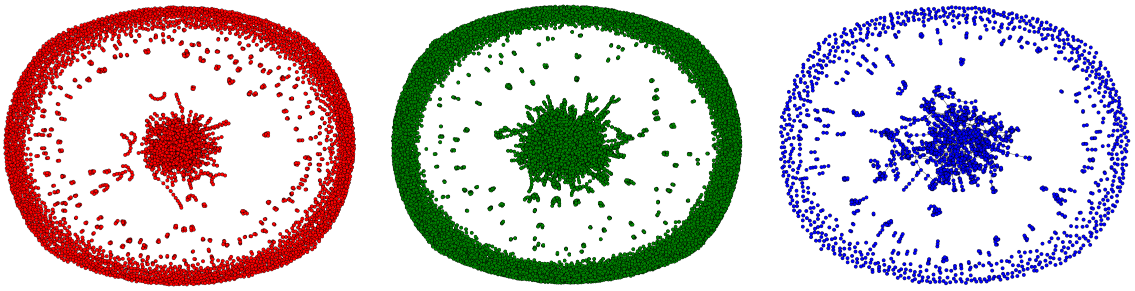

(a) EC-EC interaction. (b) Compound-Compound interaction. (c) Pathway-Pathway interaction.

Figure 2: Network visualization of EC, compound and pathway nodes from MetaCyc. As it can be observed that compounds and ECs comprise the largest part of MetaCyc.

#### 4 Dataset Characteristics

The datasets employed in this work are: i)- golden tier 1 dataset, composed of six databases, retrieved from biocyc website: *EcoCyc (v21)*, *HumanCyc (v19.5)*, *AraCyc (v18.5)*, *YeastCyc (v19.5)*, *LeishCyc (v19.5)*, and *TrypanoCyc (v18.5)*, and are refined to include only those pathways that cross-intersect with the *MetaCyc* database (v21) [1]; and ii)- BioCyc (v20.5 tier 2 & 3) [2], which consists of 9255 PGDBs (Pathway/Genome Databases) with 1463 distinct pathway labels and is constructed using the Pathway Tools software [6]. The detailed characteristics of the datasets are summarized in Table 1. For each dataset $\mathcal{S}$, we use $|\mathcal{S}|$ and $\mathrm{L}(\mathcal{S})$ to represent the number of instances and pathway labels, respectively. In addition, we also present some characteristics of the multi-label datasets, which are denoted as:

1. Label cardinality ($\mathrm{LCard}(\mathcal{S}) = \frac{1}{n}\sum_{i=1}^{i=n}\sum_{j=1}^{j=t}\mathbb{I}[\mathbf{Y}_{i,j} \neq -1]$), where $\mathbb{I}$ is an indicator function. It denotes the average number of pathways in $\mathcal{S}$.
2. Label density ($\mathrm{LDen}(\mathcal{S}) = \frac{LCard(\mathcal{S})}{L(\mathcal{S})}$). This is simply obtained through normalizing $\mathrm{LCard}(\mathcal{S})$ by the number of total pathways in $\mathcal{S}$.
3. Distinct label sets ($\mathrm{DL}(\mathcal{S})$). This notation indicates the number of distinct pathways in $\mathcal{S}$.
4. Proportion of distinct label sets ($\mathrm{PDL}(\mathcal{S}) = \frac{DL(\mathcal{S})}{|\mathcal{S}|}$). It represents the normalized version of $\mathrm{DL}(\mathcal{S})$, and is obtained by dividing DL(.) with the number of instances in $\mathcal{S}$.

The notations $\mathrm{R}(\mathcal{S})$, $\mathrm{RCard}(\mathcal{S})$, $\mathrm{RDen}(\mathcal{S})$, $\mathrm{DR}(\mathcal{S})$, and $\mathrm{PDR}(\mathcal{S})$ have similar meanings as before but for the enzymatic reactions $\mathcal{E}$ in $\mathcal{S}$. Finally, $\mathrm{PLR}(\mathcal{S})$ represents a ratio of $\mathrm{L}(\mathcal{S})$ to $\mathrm{R}(\mathcal{S})$.

#### 5 Metabolic Pathway Prediction

For this case study, we report the performance of mlLGPR (elastic-net) with concatenated features learned from cm2v on 6 benchmark datasets, as described in Section 4. Table 2 shows the comparative performances of mlLGPR+cm2v against other pathway prediction algorithms using four evaluation metrics: *Hamming loss*, *Micro precision*, *Micro recall*, and *Micro F1 score*.

#### 6 Scalability

Here, we analyze training times (after 3 epochs) under full+uec MetaCyc, where node2vec (n2v), in Fig 5, is observed to be scalable to thousands of nodes, without requiring prior knowledge on meta-paths, while metapath2vec++ (cm2v) and RUST-norm (crt) and are less likely to scale on a large graph.

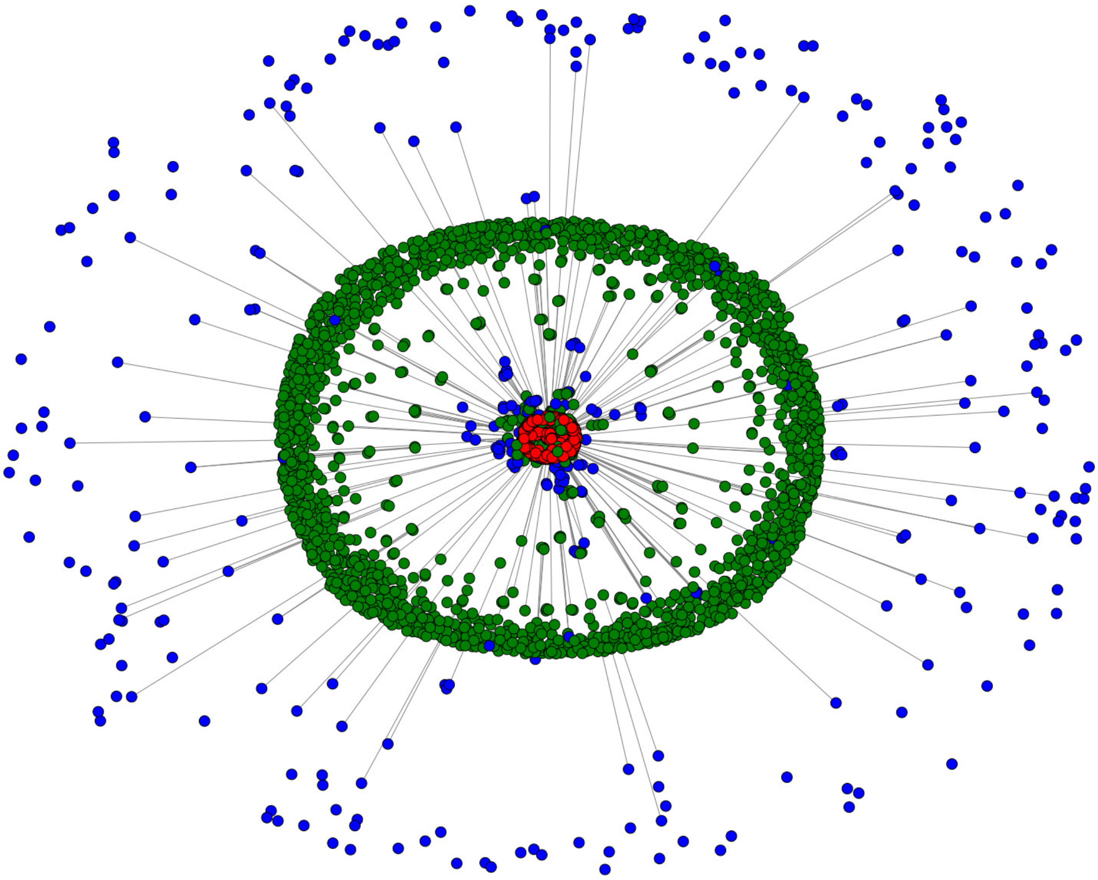

Figure 3: Network visualization of randomly selected 3000 nodes from MetaCyc. The colors corresponds to: i)- red for ECs, ii)- green for compounds, and iii)- blue to indicate pathways.

| Dataset | $\lvert\mathcal{S}\rvert$ | L($\mathcal{S}$) | LCard($\mathcal{S}$) | LDen($\mathcal{S}$) | DL($\mathcal{S}$) | PDL($\mathcal{S}$) | R($\mathcal{S}$) | RCard($\mathcal{S}$) | RDen($\mathcal{S}$) | DR($\mathcal{S}$) | PDR($\mathcal{S}$) | PLR($\mathcal{S}$) | Domain |
|---|---|---|---|---|---|---|---|---|---|---|---|---|---|
| Synset-2 | 15000 | 6806262 | 453.7508 | 0.00007 | 2526 | 0.1684 | 34006386 | 2267.0924 | 0.00007 | 3650 | 0.2433 | 0.2001 | Synthetically generated (corrupted) |
| EcoCyc | 1 | 307 | 307 | 1 | 307 | 307 | 1134 | 1134 | 1 | 719 | 719 | 0.2707 | Escherichia coli K-12 substr.MG1655 |
| HumanCyc | 1 | 279 | 279 | 1 | 279 | 279 | 1177 | 1177 | 1 | 693 | 693 | 0.2370 | Homo sapiens |
| AraCyc | 1 | 510 | 510 | 1 | 510 | 510 | 2182 | 2182 | 1 | 1034 | 1034 | 0.2337 | Arabidopsis thaliana |
| YeastCyc | 1 | 229 | 229 | 1 | 229 | 229 | 966 | 966 | 1 | 544 | 544 | 0.2371 | Saccharomyces cerevisiae |
| LeishCyc | 1 | 87 | 87 | 1 | 87 | 87 | 363 | 363 | 1 | 292 | 292 | 0.2397 | Leishmania major Friedlin |
| TrypanoCyc | 1 | 175 | 175 | 1 | 175 | 175 | 743 | 743 | 1 | 512 | 512 | 0.2355 | Trypanosoma brucei |
| BioCyc | 9255 | 1804003 | 194.9220 | 0.0001 | 1463 | 0.1581 | 8848714 | 956.1009 | 0.0001 | 2705 | 0.2923 | 0.2039 | BioCyc version 20.5 (tier 2 & 3) |

Table 1: **Characteristics of the experimental datasets.** The notations $\lvert\mathcal{S}\rvert$, L($\mathcal{S}$), LCard($\mathcal{S}$), LDen($\mathcal{S}$), DL($\mathcal{S}$), and PDL($\mathcal{S}$) represent number of instances, number of pathway labels, pathway labels cardinality, pathway labels density, distinct pathway labels set, and proportion of distinct pathway labels set for $\mathcal{S}$, respectively. The notations R($\mathcal{S}$), RCard($\mathcal{S}$), RDen($\mathcal{S}$), DR($\mathcal{S}$), and PDR($\mathcal{S}$) have similar meanings as before but for the enzymatic reactions $\mathcal{E}$ in $\mathcal{S}$. PLR($\mathcal{S}$) represents a ratio of L($\mathcal{S}$) to R($\mathcal{S}$). The last column denotes the domain of $\mathcal{S}$.

### 7 Similarity Search

We conducted cosine similarity search to determine the distance between the query pathway and the rest of pathways using metapath2vec++ (any other method discussed in this paper can be employed). For this, we selected a total of 21 nitrogen metabolic pathways. For illustration purposes,

| Methods | Hamming Loss ↓ | | | | | |
|---|---|---|---|---|---|---|
| | EcoCyc | HumanCyc | AraCyc | YeastCyc | LeishCyc | TrypanoCyc |
| PathoLogic | 0.0610 | **0.0633** | 0.1188 | **0.0424** | **0.0368** | **0.0424** |
| MinPath | 0.2257 | 0.2530 | 0.3266 | 0.2482 | 0.1615 | 0.2561 |
| mlLGPR | 0.0804 | **0.0633** | **0.1069** | 0.0550 | 0.0380 | 0.0590 |
| mlLGPR+n2v | 0.0558 | 0.1021 | 0.1706 | 0.0768 | 0.0424 | 0.0883 |
| mlLGPR+m2v | 0.0558 | 0.0998 | 0.1742 | 0.0740 | <u>0.0412</u> | 0.0926 |
| mlLGPR+cm2v | 0.0586 | 0.1041 | 0.1742 | 0.0744 | 0.0420 | 0.0867 |
| mlLGPR+jt | 0.0550 | 0.1041 | 0.1738 | <u>0.0724</u> | 0.0459 | 0.0895 |
| mlLGPR+rt | 0.0554 | <u>0.0990</u> | 0.1746 | 0.0752 | 0.0428 | <u>0.0855</u> |
| mlLGPR+crt | <u>**0.0542**</u> | 0.1017 | <u>0.1615</u> | 0.0760 | 0.0439 | <u>0.0855</u> |

| Methods | Micro Precision Score ↑ | | | | | |
|---|---|---|---|---|---|---|
| | EcoCyc | HumanCyc | AraCyc | YeastCyc | LeishCyc | TrypanoCyc |
| PathoLogic | **0.7230** | **0.6695** | 0.7011 | **0.7194** | **0.4803** | **0.5480** |
| MinPath | 0.3490 | 0.3004 | 0.3806 | 0.2675 | 0.1758 | 0.2129 |
| mlLGPR | 0.6187 | 0.6686 | 0.7372 | 0.6480 | 0.4731 | 0.5455 |
| mlLGPR+n2v | 0.7923 | 0.5745 | 0.6965 | 0.6446 | 0.4153 | 0.3974 |
| mlLGPR+m2v | 0.7862 | <u>0.6015</u> | 0.6786 | 0.6750 | <u>0.4261</u> | 0.3745 |
| mlLGPR+cm2v | 0.7770 | 0.5556 | 0.6620 | 0.6723 | 0.4159 | 0.4076 |
| mlLGPR+jt | 0.7979 | 0.5556 | 0.6732 | <u>0.6949</u> | 0.3840 | 0.3924 |
| mlLGPR+rt | 0.7889 | 0.6014 | 0.6635 | 0.6560 | 0.4146 | <u>0.4113</u> |
| mlLGPR+crt | <u>**0.7993**</u> | 0.5873 | <u>**0.7898**</u> | 0.6581 | 0.3983 | 0.4105 |

| Methods | Micro Recall Score ↑ | | | | | |
|---|---|---|---|---|---|---|
| | EcoCyc | HumanCyc | AraCyc | YeastCyc | LeishCyc | TrypanoCyc |
| PathoLogic | 0.8078 | 0.8423 | 0.7176 | 0.8734 | 0.8391 | 0.7829 |
| MinPath | **0.9902** | **0.9713** | **0.9843** | **1.0000** | **1.0000** | **1.0000** |
| mlLGPR | 0.8827 | 0.8459 | 0.7314 | 0.8603 | 0.9080 | 0.8914 |
| mlLGPR+n2v | 0.7329 | 0.2903 | 0.2745 | 0.3406 | 0.5632 | 0.5314 |
| mlLGPR+m2v | <u>0.7427</u> | 0.2867 | 0.2608 | 0.3537 | 0.5632 | 0.5029 |
| mlLGPR+cm2v | 0.7264 | 0.2867 | <u>0.2804</u> | 0.3493 | 0.5402 | <u>0.5543</u> |
| mlLGPR+jt | 0.7329 | 0.2867 | 0.2706 | <u>0.3581</u> | 0.5517 | 0.5314 |
| mlLGPR+rt | <u>0.7427</u> | <u>0.3082</u> | 0.2745 | <u>0.3581</u> | <u>0.5862</u> | 0.5429 |
| mlLGPR+crt | 0.7394 | 0.2652 | 0.2725 | 0.3362 | 0.5402 | 0.5371 |

| Methods | Micro F1 Score ↑ | | | | | |
|---|---|---|---|---|---|---|
| | EcoCyc | HumanCyc | AraCyc | YeastCyc | LeishCyc | TrypanoCyc |
| PathoLogic | 0.7631 | 0.7460 | 0.7093 | **0.7890** | 0.6109 | 0.6447 |
| MinPath | 0.5161 | 0.4589 | 0.5489 | 0.4221 | 0.2990 | 0.3511 |
| mlLGPR | 0.7275 | **0.7468** | **0.7343** | 0.7392 | **0.6220** | **0.6768** |
| mlLGPR+n2v | 0.7614 | 0.3857 | 0.3938 | 0.4457 | 0.4780 | 0.4548 |
| mlLGPR+m2v | 0.7638 | 0.3883 | 0.3768 | 0.4642 | 0.4851 | 0.4293 |
| mlLGPR+cm2v | 0.7508 | 0.3783 | 0.3939 | 0.4598 | 0.4700 | <u>0.4697</u> |
| mlLGPR+jt | 0.7640 | 0.3783 | 0.3860 | <u>0.4726</u> | 0.4528 | 0.4515 |
| mlLGPR+rt | 0.7651 | <u>0.4076</u> | 0.3883 | 0.4633 | <u>0.4857</u> | 0.4680 |
| mlLGPR+crt | <u>**0.7682**</u> | 0.3654 | <u>0.4052</u> | 0.4451 | 0.4585 | 0.4653 |

Table 2: **Predictive performance of each comparing algorithm on 6 benchmark datasets.** For each performance metric, '↓' indicates the smaller score is better while '↑' indicates the higher score is better.

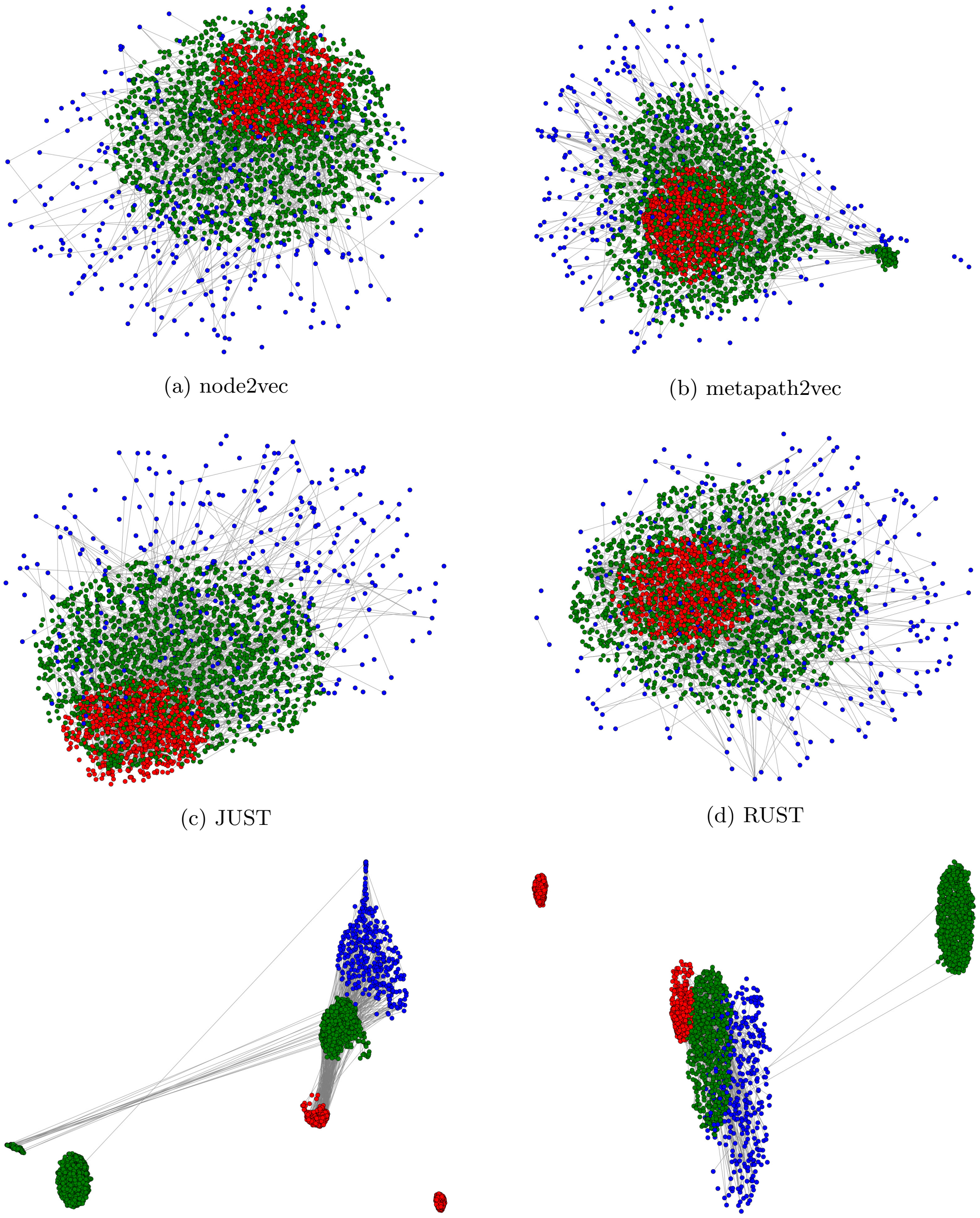

(a) node2vec (b) metapath2vec

(c) JUST (d) RUST

(e) metapath2vec++ (f) RUST-norm

Figure 4: 2D UMAP projections of 3000 nodes. The colors corresponds to: i)- red for ECs, ii)- green for compounds, and iii)- blue to indicate pathways. Obviously, metapath2vec++ and RUST-norm in Figs 4e and 4f have clear distinct boundaries in contrast to node2vec, metapath2vec, JUST, and RUST in Figs 4a, 4b, 4c, and 4d. The islands corresponds nodes with links lower than 2.

Table 3 lists only the top 5 results for querying the 7 pathway ids. One can observe that for the query id "DENITRIFICATION-PWY" (nitrate reduction I (denitrification)), for example, cm2v returns pathways that are variants to it, such as "PWY-5674" (nitrate reduction IV (dissimilatory)) and "PWY-5675" (nitrate reduction V (assimilatory)). Similar results can be also recovered when querying other pathways (or ECs).

### References

[1] Ron Caspi, Richard Billington, Luciana Ferrer, Hartmut Foerster, Carol A. Fulcher, Ingrid M. Keseler, Anamika Kothari, Markus Krummenacker, Mario Latendresse, Lukas A. Mueller,

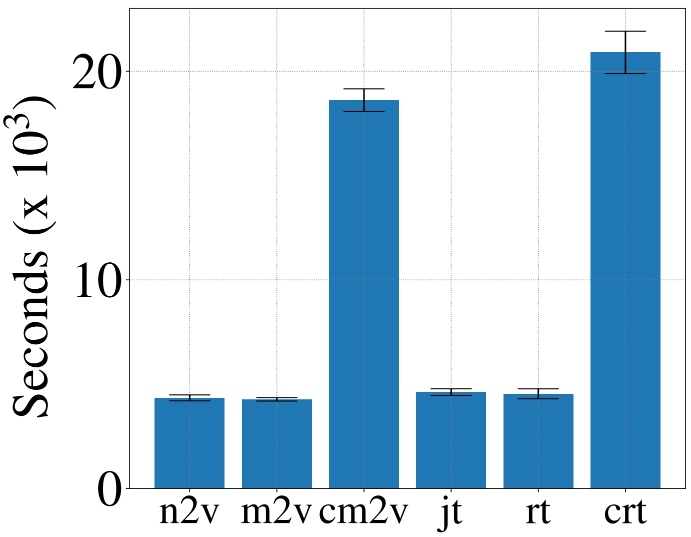


Figure 5: Scalability measured in seconds ($\times 10^3$) under uec+full configuration.

| Rank | PWY-6964 | AMMOXID-PWY | P303-PWY | PWY-2242 |
|---|---|---|---|---|
| 1 | GLUTAMINDEG-PWY | PWY-5366 | PWY-7058 | PWY-1269 |
| 2 | PWY-7672 | PWY-6014 | PWY-862 | AMMOXID-PWY |
| 3 | GLUTAMINEFUM-PWY | PWY-5373 | PWY-7562 | HEMESYN2-PWY |
| 4 | CITRULBIO-PWY | PWY-7802 | PWY-5789 | PWY-6557 |
| 5 | PWY-5675 | PWY-6837 | PWY-6310 | PWY-6873 |
| Rank | DENITRIFICATION-PWY | PWY490-3 | PWY-1264 | TAURINEDEG-PWY |
| 1 | PWY-5674 | PWY-5675 | PWY-5944 | PWY-5046 |
| 2 | PWY-5675 | PWY-381 | TAURINEDEG-PWY | PWY-6043 |
| 3 | PWYQT-4471 | PWY-6840 | PWY-5844 | PWY-282 |
| 4 | PWY-7405 | TRNA-CHARGING-PWY | PWY-6423 | PWY-6388 |
| 5 | PWY-6275 | PWY-6945 | PWY-1263 | PWY-7833 |

Table 3: Top 5 Pathway IDs for nitrogen metabolism.


Quang Ong, Suzanne Paley, Pallavi Subhraveti, Daniel S. Weaver, and Peter D. Karp. The metacyc database of metabolic pathways and enzymes and the biocyc collection of pathway/genome databases. *Nucleic Acids Research*, 44(D1):D471–D480, 2016.

[2] Ron Caspi, Richard Billington, Hartmut Foerster, Carol A Fulcher, Ingrid Keseler, Anamika Kothari, Markus Krummenacker, Mario Latendresse, Lukas A Mueller, Quang Ong, et al. Biocyc: Online resource for genome and metabolic pathway analysis. *The FASEB Journal*, 30(1 Supplement):lb192–lb192, 2016.

[3] Yuxiao Dong, Nitesh V Chawla, and Ananthram Swami. metapath2vec: Scalable representation learning for heterogeneous networks. In *Proceedings of the 23rd ACM SIGKDD International Conference on Knowledge Discovery and Data Mining*, pp. 135–144. ACM, 2017.

[4] Aditya Grover and Jure Leskovec. node2vec: Scalable feature learning for networks. In *Proceedings of the 22nd ACM SIGKDD international conference on Knowledge discovery and data mining*, pp. 855–864. ACM, 2016.

[5] Rana Hussein, Dingqi Yang, and Philippe Cudré-Mauroux. Are meta-paths necessary?: Revisiting heterogeneous graph embeddings. In *Proceedings of the 27th ACM International Conference on Information and Knowledge Management*, pp. 437–446. ACM, 2018.

[6] Peter D Karp, Mario Latendresse, Suzanne M Paley, Markus Krummenacker, Quang D Ong, Richard Billington, Anamika Kothari, Daniel Weaver, Thomas Lee, Pallavi Subhraveti, et al. Pathway tools version 19.0 update: software for pathway/genome informatics and systems biology. *Briefings in bioinformatics*, 17(5):877–890, 2016.

[7] Bryan Perozzi, Rami Al-Rfou, and Steven Skiena. Deepwalk: Online learning of social representations. In *Proceedings of the 20th ACM SIGKDD international conference on Knowledge discovery and data mining*, pp. 701–710. ACM, 2014.
